# Brittleness in model selection analysis of single neuron firing rates

**DOI:** 10.1101/430710

**Authors:** Chandramouli Chandrasekaran, Joana Soldado-Magraner, Diogo Peixoto, William T. Newsome, Krishna V. Shenoy, Maneesh Sahani

**Author notes:** Corresponding Authors: CC, KVS, MS. denotes equal contribution (Krishna V. Shenoy, Maneesh Sahani).

## Abstract

Models of complex heterogeneous systems like the brain are inescapably incomplete, and thus always falsified with enough data. As neural data grow in volume and complexity, absolute measures of adequacy are being replaced by model selection methods that rank the relative accuracy of competing theories. Selection still depends on incomplete mathematical instantiations, but the implicit expectation is that ranking is robust to their details. Here we highlight a contrary finding of “brittleness,” where data matching one theory conceptually are ranked closer to an instance of another. In particular, selection between recent models of decision making is conceptually misleading when data are simulated with minor distributional mismatch, with mixed secondary signals, or with non-stationary parameters; and decision-related responses in macaque cortex show features suggesting that these effects may impact empirical results. We conclude with recommendations to mitigate such brittleness when using model selection to study neural signals.

## Introduction

Sciences that deal with heterogeneous complex systems—including ecology, economics, finance, medicine and systems neuroscience—encounter a particular challenge when seeking to validate theories with data. The classical scientific method, exemplified in fundamental physics, requires careful quantitative experimental measurements to corroborate precise theoretical predictions within the margin of observational error (e.g., ATLAS Collaboration, 2012; Schneider, 1992; Renn et al., 1997). But this is rarely possible in a complex system: no tractable formal theory will fully account for every complexity, and so measurements are influenced by factors unmodeled and typically uncontrollable (Focardi et al., 2012). Thus, theories in these fields are most often loosely-defined models or working hypotheses that may capture the essential trends of empirical phenomena but inevitably simplify the mathematical forms of empirical relationships, and the shape and source of empirical variability.

At the most basic level, progress in these sciences can be made using experiments which simply test for the existence of a hypothesized relationship in the data. For example, do primary visual cortical neurons modulate their firing in response to the orientation of a contrast edge (Hubel and Wiesel, 1959)? Other studies seek an ordering of effects: is one drug treatment more effective than another (Kane et al., 1988)? For these types of questions, the classical statistical hypothesis testing framework (Neyman and Pearson, 1966) can reject null hypotheses of independence or of equivalent efficacy. However, the approach does not easily extend to testing models: for instance, are the dynamics of grating-evoked responses consistent with a model where tuning curves are shaped by strong recurrent interaction (Ringach et al., 1997)? In such cases, the null hypothesis that should be tested using observational data is not obvious, and direct evaluation of the validity of a particular model of recurrent interaction inevitably shorn of the fine details of the circuit (e.g., Ben-Yishai et al., 1995), will always lead to rejection in the face of sufficient data. In some cases, a suitably-designed intervention may help to address the issue (Lien and Scanziani, 2013; Reinhold et al., 2015) but for many models, interventions are both conceptually and technologically intractable.

The unsuitability of Neyman-Pearson hypothesis testing to validate models of complex systems has led many to argue for the use of *model selection* in its place (Aho et al., 2014; Anderson and Burnham, 2002; Raftery, 1995). This family of approaches, which includes cross-validation (Gelman et al., 2014), a variety of information criteria (Akaike, 1974; Hannan and Quinn, 1979; Akaike, 1998; Aho et al., 2014; Gelman et al., 2014; Spiegelhalter et al., 2002, 2014), and Bayesian model evidence or Bayes factors (Gelman et al., 2014; Kass and Raftery, 1995), compares two or more different models—representative of two or more working hypotheses—to select the model that provides a better account for a set of observations. Recent years have seen burgeoning interest in applying model selection in neuroscience, both for the firing patterns of single neurons (Bollimunta et al., 2012; Latimer et al., 2015b,a, 2016, 2017; Rossant et al., 2011), as well as for functional imaging and encephalographic signals (Durstewitz et al., 2016; Linderman and Gershman, 2017; Marreiros et al., 2010a,b; Mars et al., 2012). While broadly supportive, previous authors have highlighted potential pitfalls and challenges to the proper application of model selection approaches in neuroscience and other fields (Aho et al., 2014; Anderson and Burnham, 2002; Churchland and Kiani, 2016; Mars et al., 2012). Much of this concern has focused on the properties and failures of particular selection criteria, especially in situations where one or the other of the models is assumed to be correct or closest in a chosen sense to the data (Gelman et al., 2014).

In this viewpoint, we highlight a deeper, conceptual, challenge to the application and interpretation of model selection; one that emerges from a common lack of robustness, which we term “brittleness,” of selection results in the face of small and apparently tangential deviations between models and data. We encountered this brittleness as we sought to expand on recent results using model selection (Latimer et al., 2015b, 2017) and so present our findings in the context of that study, but note that qualitatively similar issues have arisen in other fields (e.g., ecology; Anderson and Burnham, 2002) and so the issue is very likely to be pervasive, although it appears to be still underappreciated in neuroscience and allied fields.

The brittleness arises from the need to translate loose theoretical working hypotheses, framed at a conceptual or intuitive level, into precise mathematical models for testing. Consider, for instance, the question of whether a neuron signals a decision by an abrupt transition in firing at one point in a trial (a “step”) or alternatively reflects the gradual accumulation of evidence by a graded change in rate across the entire trial duration (a “ramp”) (Shadlen and Newsome, 1996; Okamoto et al., 2007; Bollimunta et al., 2012; Latimer et al., 2015b). These are conceptually distinct theories of single-neuron dynamics during decision making, but as they stand they are too loosely defined for model selection. The broad classes they represent must first be reduced to specific mathematical instances. Latimer and colleagues chose a doubly stochastic “step” model in which a single transition between fixed spike rates happened at a time and in a direction that depended on sensory evidence, with spike times distributed according to a Poisson process given the stepped rate. Their doubly stochastic “ramp” model was based on the drift-diffusion model (DDM) for decision-making: a latent variable integrated sensory evidence over time until it reached a threshold after which it remained constant. The sensory evidence was assumed to be drawn from a noisy Gaussian distribution. The latent variable was mapped to firing through a soft-threshold non-linearity to yield a predicted rate, with spike times once again drawn from a Poisson process (Latimer et al., 2015b).

These are both plausible and simple representative candidate models for steps and ramps, but it cannot reasonably be expected that real neurons’ firing will be described exactly by either one. Nonetheless, many model selection criteria seek to identify the model that is closest in some sense (typically according to a probability divergence, such as the Kullback-Leibler divergence), or following Box (1976), the most “useful” in predictive terms (Aho et al., 2014; Box, 1976; Burnham and Anderson, 2003). The presumption is that this sort of criterion will be robust, in that data generated by any process that falls into one of the broad conceptual hypothesis classes will be closer to the particular mathematical model used to instantiate that class; or, equivalently, that that mathematical instantiation will be the more useful. If a neuron’s firing steps, then model selection should prefer the Poisson change-point step process even if the true step statistics differ from those assumed. Conversely, if the true change in firing is gradual, like a ramp, then the drift-based model should be more predictive even if the true ramp follows a different distribution.

Our findings show that this robustness does not always hold. Synthetic data that are generated with small deviations from the drift-diffusion model, such as underdispersion of spike counts relative to Poisson or incorporation of a systematic stimulus-independent modulation of the rate, become reliably associated with the “step” model even though no firing-rate step is introduced. More worryingly, data are similarly misassigned even if they are simulated from the *exact* form of drift-diffusion model used for selection, but with different settings of the parameters in different subsets. Examining neuronal data from the lateral intraparietal cortex (Rorie et al., 2010) and dorsal premotor cortex (Chandrasekaran et al., 2017), we find relationships between firing statistics and model selection results, as well as disparities in trial-by-trial contributions to the selection results, that are consistent with these potential forms of misassignment. Thus selection results based on these two specific mathematical models can at best provide only weak evidence to decide between the “step” and “ramp” conceptual classes that they represent.

The conceptual difficulty in the interpretation of model selection applied to complex heterogeneous systems that our results highlight is very likely to apply in other settings in neuroscience and beyond. Nonetheless, progress in understanding such systems will ultimately depend on deploying these methods with care. We offer some suggestions in the Discussion to help in the mitigation of the attendant challenges.

A preliminary version of these results was reported previously (Chandrasekaran et al., Society For Neuroscience Annual Meeting, 2016).

## Results

We evaluated the robustness of model selection using the same selection criterion and the same specific mathematical models of “step” and “ramp” decision making as adopted by Latimer et al. (2015b). We considered three ways in which recorded neural data might plausibly differ from both of the mathematical models being compared, while in each case remaining broadly coherent with the ramp conceptual framework. Each variation was motivated by either a commonly observed feature of neural activity, or a theoretical consideration regarding decision-making circuitry. In each case, we first simulated data from the variant ramp model, evaluating the results of model selection using Latimer and colleagues’ approach. We then examined decision-related neural data collected in two different brain regions for signs that the effects we noted in the simulations might also shape the results of empirical model selection.

### Neural Data

We used two data sets, drawn respectively from the lateral intraparietal area (LIP) and the dorsal aspect of the pre-motor cortex (PMd) in macaque monkeys making binary-outcome decisions about sensory stimuli.

#### LIP Data

The LIP data were recorded by Rorie and collaborators (Rorie et al., 2010) from two monkeys performing an oculomotor random-dot motion discrimination task (Supp. Fig. 1A). We used data from 81 LIP neurons recorded across 4 reward conditions, providing us with a pool of 324 (81×4) pseudo-independent recordings. As suggested in the Latimer et al. (2015b) study, we considered a recording for analysis only if it had sufficient choice selectivity (defined as having a signal-detection theory sensitivity (*d*′) of 0.5 or greater). This criterion selected 117 recordings. The firing rates and other response properties of these recordings are consistent with the many reports of decision-related activity in LIP (e.g., Shadlen and Newsome, 2001) and other brain areas (Ding and Gold, 2012; Hanks et al., 2015). LIP neurons had strong firing rates for the preferred (PREF) saccadic choice trials and modest decreases or flat firing rates for the nonpreferred (NONPREF) saccadic choices.

#### PMd Data

The PMd data comprised 806 units recorded from two monkeys while they performed a color checkerboard reaction time discrimination task, reporting their decision with a self-timed arm movement (Supp. Figs. 1B-D). We have previously shown that trial-averaged firing rates of a subpopulation of these PMd units increase gradually during the trial, at a rate that covaries systematically with choice, sensory evidence, and reaction time (Chandrasekaran et al., 2017). These are features described in many other studies of decision-related activity in various cortical and subcortical areas (e.g., Roitman and Shadlen, 2002; Shadlen and Kiani, 2013; Thura and Cisek, 2014) and consistent with properties expected of a candidate decision variable (Gold and Shadlen, 2007). In the current study, we examined 429 units that showed the expected hallmarks of decision-related activity (Shadlen and Kiani, 2013). The average firing rate of the 429 units organized by stimulus coherence and choice, and reaction time and choice, are shown in Supp. Figs. 2A, B. Other units either decreased their activity or only responded just before movement initiation and were inconsistent with a candidate decision variable. We again enforced the criterion that *d*′ ≥ 0.5, selecting a total of 311 units.

### Model fitting and DIC

We used the computer code developed and generously shared by Latimer and colleagues to perform a Bayesian fit of both the DDM and the step model to each neuron in both data sets, obtaining Monte-Carlo samples of parameters that were consistent *a posteriori* with the measured spike counts (10 ms bins). These “posterior” samples allowed us to evaluate the difference in the deviance information criterion (DIC) between the two models (Spiegelhalter et al., 2002). DIC is a model selection metric that combines a goodness of fit term (measured using the deviance or scaled negative log-likelihood relative to a baseline) with a penalty for model complexity. Higher values of DIC indicate a poorer description of the data by the model. The difference between the DIC for the DDM and the step model, the “DIC score,” provides an estimate of the log-ratio of deviances penalised by a measure of relative model complexity, and can be used to rank the support offered by the data for the different models. As per Latimer et al. (2015b), negative DIC scores suggest that the data are better described by a DDM; conversely, positive DIC scores signal data that appear closer to the step model.

Although the precise form of penalty assumed by DIC has been criticised (Spiegelhalter et al., 2014), it remains widely used. We adopted it here to maintain consistency with the methods of Latimer et al. (2015b), and because our goal was to address a general point about model selection that is broadly independent of the criterion used. Further experiments, not shown, revealed comparable results using alternative model selection criteria.

The total deviance and complexity penalty are both obtained by summing contributions from individual trials, and so the magnitude of DIC tends to increase with trial count. Thus, although we found larger magnitudes of DIC score in PMd than in either our own LIP data, or than reported by Latimer et al. (2015b) in theirs, the disparity was very likely to have arisen at least partly from the significantly larger number of trials we had available from PMd. Furthermore, as both expected value and variance of DIC scores increase with trial count, and trial length, numerical comparison of scores across studies is difficult even for similar neuronal populations, as is the choice of a single absolute threshold value for “significance” (Murtaugh, 2014). We therefore interpreted any DIC score > 0 as evidence in support of the step model, and scores < 0 as evidence for the DDM.

### 1. Brittleness to violation of parametric assumptions

Model selection is designed to evaluate parametric statistical models, which assign a probability of occurrence to each possible observation. Thus, the specific models being evaluated must incorporate assumptions, explicit or implicit, about the forms of the relevant probability distributions. Even where models are specified in hierarchical form, so that the parameters that most directly govern the probabilities themselves vary under the control of higher-level parameters, the ranges allowed and the parametric forms that they govern must still be assumed. This raises the question of how robust model selection proves to be when data are generated from a process that broadly follows the form of one of the candidate models, but differs from it—and potentially also from the competitor model—by an apparently minor parametric choice.

We considered this question in the context of the step and ramp models. Both models assumed that spike counts followed a Poisson distribution around a mean firing rate that itself varied from trial to trial. Thus, they both predicted that the total variance in the observed spike count in each bin should be overdispersed relative to Poisson, with a Fano factor greater than 1. Moreover, the DDM also predicts that the Fano factor should often increase over time, as long as neither the saturating bound nor the rectifying floor is reached routinely (Supp. Fig. 3).

In principle, biophysical processes such as firing refractoriness and adaptation might lead to sub-Poisson variability in firing. Indeed, underdispersion of spike counts has been reported before in LIP data (Maimon and Assad, 2009). In our LIP data, we found some neurons with super-Poisson Fano factors, according with the model assumptions (Fig. 1A). However, many other neurons had Fano factors less than or around 1 and in some cases these Fano factors also decreased over time (Figs. 1B-D). In fact, the median Fano factor in the 450 – 700 ms epoch onset after dots onset for PREF choices was 0.95 (min: 0.56, max: 1.40; sign test *H*_0_: Fano factor=1, p=2.17×10^−4^,) and 0.96 for NONPREF choices (min: 0.57, max: 1.42; sign test, *H*_0_: Fano factor=1, p=2.11×10^−5^). As this underdispersion reflected a parametric departure from both models, it offered a plausible test of robustness to such variation.

**Figure 1:**
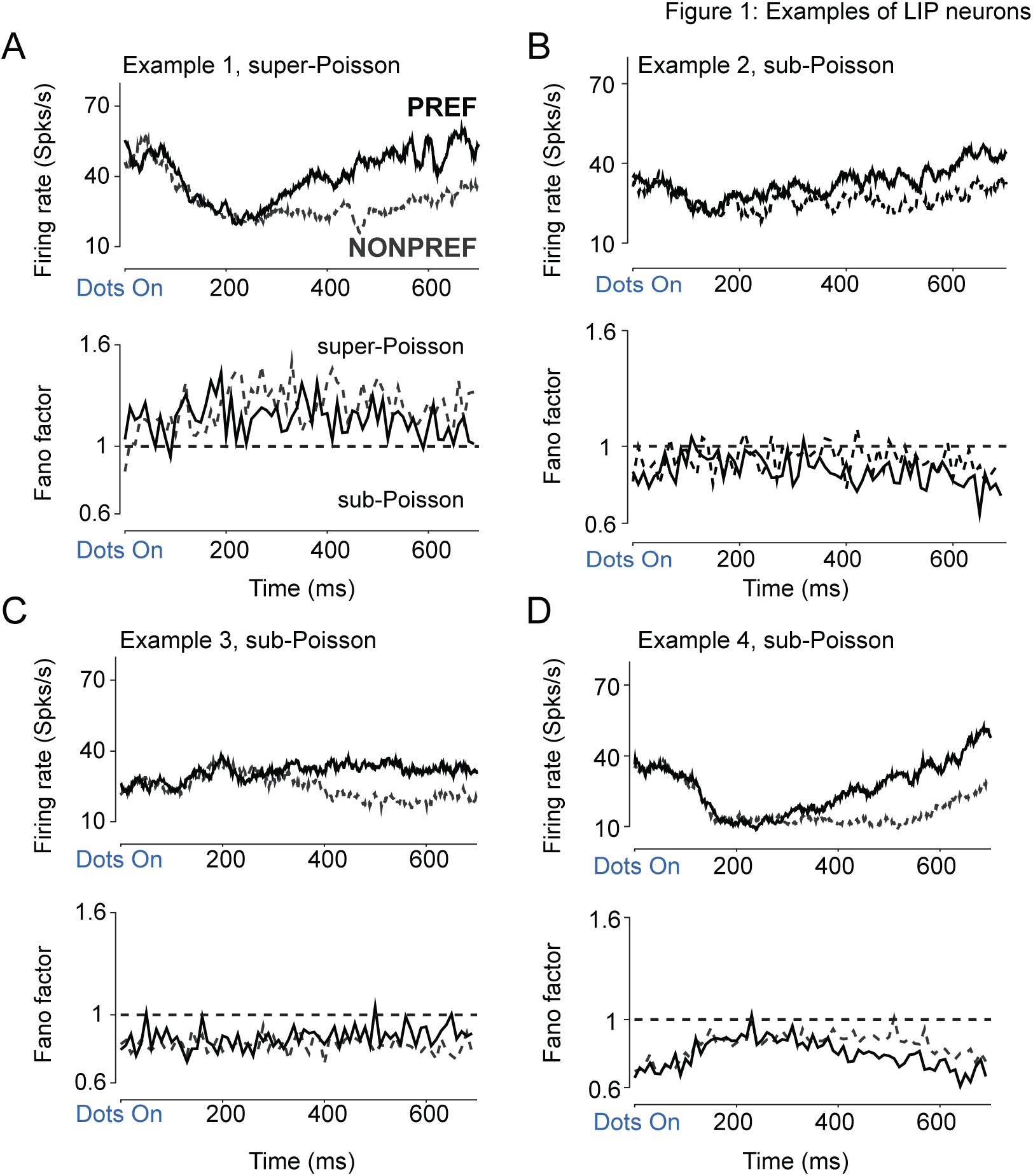
Single neurons in LIP show choice selectivity and sub-Poisson Fano factors during the decision-formation period. **A:** Example of a typical decision-related LIP neuron with robust firing rate modulation during the decision-formation period, and super-Poisson variability. **B**, **C**, **D**: Examples of LIP neurons that show modest to robust choice selectivity for the PREF direction, and sub-Poisson variability. For each neuron, the upper plot shows firing rate in spikes/s and the lower plot the time-varying Fano factor. Trials are aligned to the onset of the 500 ms-long random dot stimulus, and separated by whether the saccade was to the PREF (solid lines) or NONPREF (dashed lines) direction for each neuron. Firing rates are shown for the “LL” reward condition where each correct choice is rewarded with one drop of juice, and computed by smoothing spike trains with a 50 ms causal box car filter and then averaging over ~ 65 trials. Fano factors for the same groups of trials were estimated in non-overlapping bins of 10 ms each.

#### Simulated underdispersed DDM firing rates are assigned to the step model

We began by simulating data using the drift-diffusion latent process of the ramp model, but with spikes generated by a process (gamma-renewal) that is underdispersed relative to Poisson (see Methods and Supp. Fig. 4A). In these simulations we assumed strong sensory evidence (a non-zero drift rate) in one group of trials and no sensory evidence (zero drift rate) in the other group, corresponding to pattern seen in much of our data. While these simulations with underdispersed spike counts depart from the exact form of DDM used in model selection, they do not introduce any sort of discrete step. Thus, if model selection were robust we might expect no more than a modest lowering of confidence in the selection, perhaps reflected in a reduced magnitude of DIC score favoring the DDM. In fact, we found a reliable trend for model selection to favour the step model as an explanation of these simulated sub-Poisson DDM spike trains (e.g., Figs. 2A-C and Figs. 2D, E) whereas nearly identical simulations with DDM dynamics and Poisson spike-generation processes were robustly classified as consistent with the DDM (Fig. 2F). A similar bias was seen in simulations based on multiple levels of sensory evidence (4 conditions with 4 drift rates, Supp. Figs. 4B, C). This bias in favour of the step model for data generated by a process firmly within the ramp conceptual class demonstrates a brittleness of model selection to violations of underlying parametric assumptions, even when the violation is of an assumption common to both the models being compared.

**Figure 2:**
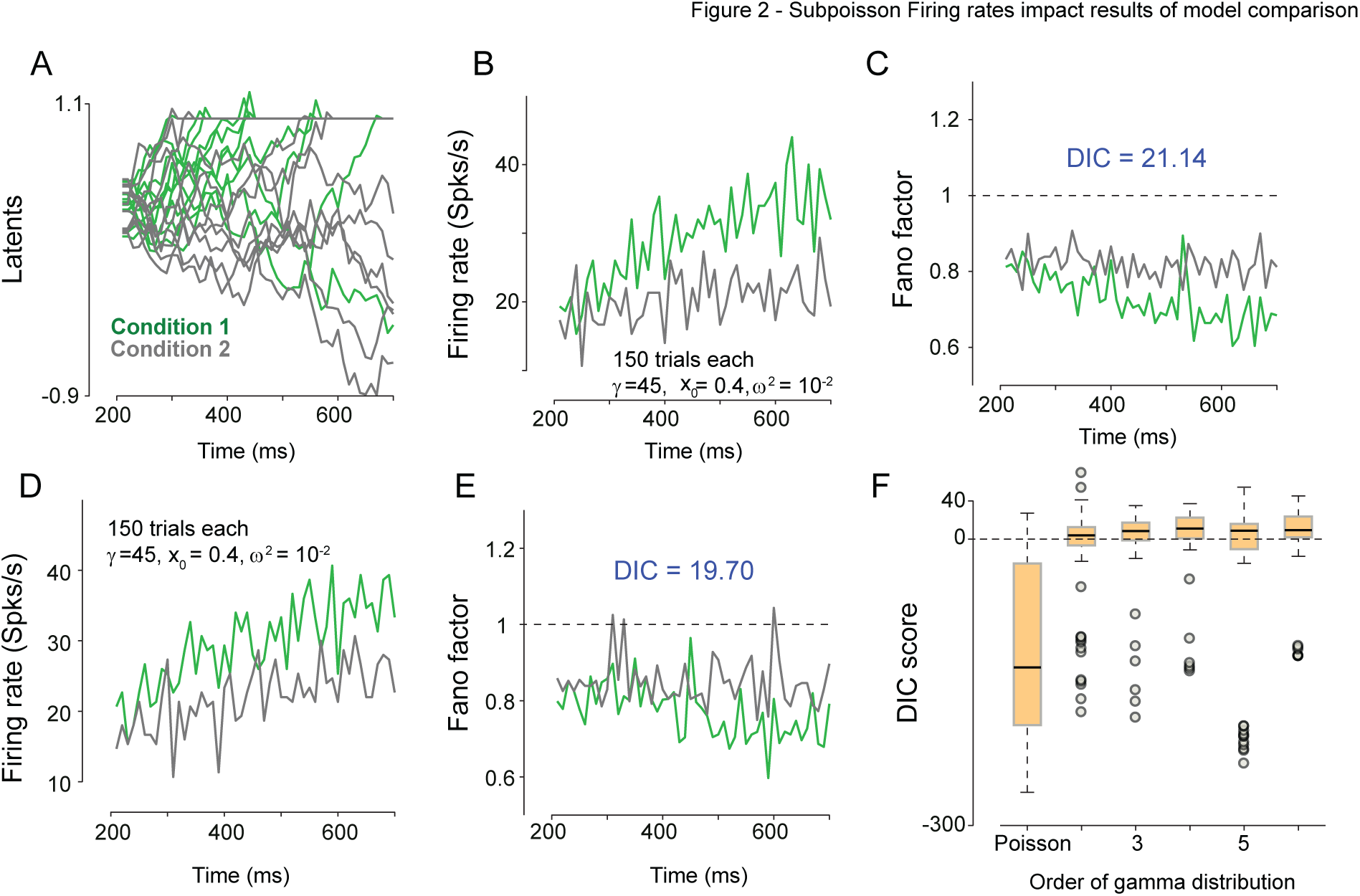
Simulated neurons with DDM dynamics and sub-Poisson spiking statistics are misleadingly classified as stepping. **A**: Dynamics of latent variable (including absorbing upper bound) for a simulated neuron from the drift diffusion model. 10 trials are shown for each condition: non-zero drift rate (condition 1, green lines) and zero drift rate (condition 2, gray lines). **B**: Trial averaged firing rates for the same simulation as in A. Firing rates are averaged over 150 trials in non-overlapping 10 ms bins. Spike trains were generated using sub-Poisson statistics with a gamma order (shape parameter) of 5. **C**: Fano factors for the simulated neuron in A, B. Fano factors were estimated using spike counts in non-overlapping 10 ms bins. A value below one shows that overall spiking statistics remain sub-Poisson even after diffusion variability across trials is included. The DIC score for this simulated neuron favors the step model even though the underlying latent dynamics corresponsed to the DDM. **D**-**E:** Firing rates (D) and Fano factor (E) for another example hypothetical neuron with sub-Poisson spiking statistics (generated with a gamma order of 3). and DDM latent dynamics but identified by model selection as consistent with the step model. **F**: Box plot of the DIC score as a function of the gamma-interval shape parameter. Higher values of this shape parameter induce increased spiking regularity and thus sub-Poisson Fano factors. DIC scores for simulated DDM neurons with Poisson statistics (shape parameter 1: no model mismatch) were correctly identified as consistent with the DDM. However, even a minor departure to a neuron with DDM dynamics and sub-Poisson spiking statistics often led many simulations to be identified as consistent with the step model. Gray dots denote outlier DIC scores either in favour of the DDM or the step model.

#### Underdispersion in data is predictive of DIC score for LIP neurons

We wondered if there might be signs of a similar dispersion-induced model selection bias in neuronal responses recorded from LIP. Unfortunately, spike-count dispersion—as varied in our simulations— could not be measured directly. Trial-to-trial variance in spike counts conflates two contributions: one from randomness in integration or stepping across trials and the second from neuronal spiking variability. Dispersion corresponds to second of these, but it cannot easily be isolated. However, we reasoned that the Fano factor on NONPREF trials, where firing rates were low and may sometimes have reflected firing at a baseline rate, might provide a indicative signal for dispersion. Thus, we asked if this Fano factor predicted DIC scores in the LIP data, either alone or in combination with additional regressors which might help to isolate dispersion more completely.

We first regressed the DIC score for each neuron against its NONPREF Fano factor in an interval 450–700 ms after dots onset, when onset-related transients should have decayed. The result was consistent with the simulation findings and the interpretation of the NONPREF variance as indicative of dispersion, with a significant regression weight between NONPREF Fano factor and DIC score for LIP neurons (*R*^2^=0.038, p=.034; *β_Fanofactor_*=−95.08±44.44 (standard error; SE); t(115)=−2.13, p=.034, see Eqn. 1).

Fano factor will be most accurate as a surrogate for dispersion when other sources of variance are small. While integration or stepping may play a role in all trials, we reasoned that their influence on variance would be greatest when they most strongly modulated firing rate, and that this modulation depth could be estimated by measuring the starting and ending firing rate during the trial. Thus, we fit an expanded multivariate regression model (see Methods, Eqn. 2) which included the starting firing rate measured between 300–400 ms (i.e. after the initial 200 ms initial dip in the LIP response) and the ending firing rate, which was estimated at 600–700 ms after dots onset for each of the PREF and NONPREF conditions. This combined regression model predicted more variance in the DIC scores (*R*^2^=0.29, p=2.17×10^−7^). In this expanded regression, Fano factor was more strongly related to the DIC score (*β_Fanofactor_*=−163.71±42.15, t(111) = −3.88, p <.0001, partial *R*^2^ for Fano factor=0.093). The coefficient for the Fano factor remained negative suggesting that underdispersion still favoured the step model. This was consistent with the results of the simulated underdispersed DDM.

The regression coefficients for starting firing rate for PREF and NONPREF choices also predicted the DIC score (PREF: *β_Start_*=6.5±1.62, t(111)=4.00, p <0.0001, NONPREF: *β_Start_*=−5.13± 1.73, t(111)=−2.96, p <.003). We also observed a modest relationship between ending firing rate for PREF and NONPREF conditions and DIC (PREF: *β_end_*=−0.27± .80, t(111)=−0.33, p=0.73, NONPREF: *β_end_*=−2.02± 1.73, t(111)=−1.98, p= .049).

Thus, although we cannot be sure of the parametric form that underlies the real data, we found that LIP responses could be underdispersed, and that DIC scores for neurons with lower Fano factors were more likely to support the step model even though the step model also predicts overdispersion. Thus, at least some of these cases may have arisen from a systematic mismatch in dispersion with both models, rather than any true evidence for a step-like process.

### 2. Brittleness emerging from mixed responses

The behaviour of each element in a complex system is often influenced by many processes running in parallel. In neuroscience, neurons with “mixed responses” that reflect more than one aspect of a task are particularly widespread in areas of the brain concerned with cognitive functions (see for e.g., Meister et al., 2013; Rigotti et al., 2013; Mante et al., 2013; Raposo et al., 2014). We wondered how the presence of such signals extraneous to model predictions might affect model selection. In particular, we considered the case of extraneous signals which very clearly fell outside conceptual repertoire of the models being tested.

The canonical average response profile associated with the decision-making process increases or decreases monotonically from an initial value towards a bound. Both DDM and step models were designed to produce such profiles. (Note that although both up and down transitions are possible in the step model given fixed sensory evidence, the timing of transition does not depend on its sign. So combining both directions of step still yields a monotonic average.)

The PMd data we considered displayed many of the hallmarks of decision making activity (Chandrasekaran et al., 2017) and indeed firing rates of some neurons increased monotonically during the decision-formation period as expected (Fig. 3A, B). However, firing rates of other neurons undulated during the trial (Fig. 3C, D and many further examples in Supp. Fig. 5). This non-monotonicity was evident in firing rate averages for single conditions, and so it was not created by averaging over different condition-specific temporal profiles. Instead, other processes, perhaps related to preparation of the upcoming movement, appear to be mixed with the decision-making signals.

**Figure 3:**
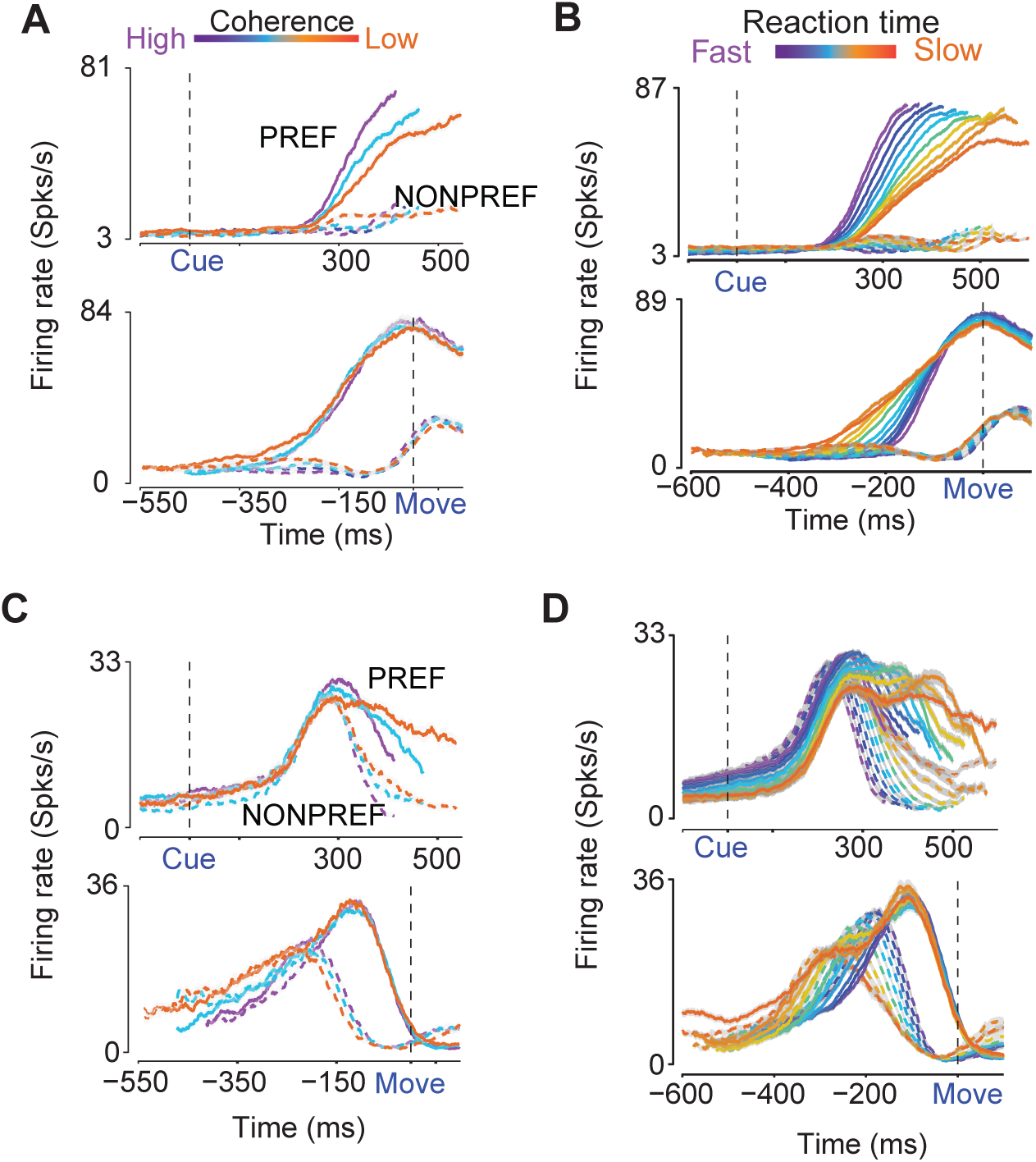
Example neuronal responses in PMd reflect both canonical decision-related activity and mixed responses. **A**: Firing rates of an example increased neuron in PMd during the decision task, sorted by coherence and arm-movement choice and aligned to checkerboard onset (CUE, top panel) or movement onset (MOVE, bottom panel). Colors label coherence conditions from easy (purple) to difficult (orange). Solid lines show movements to the PREF-direction; dashed lines show the NONPREF-direction. Firing rate traces are obtained using 75ms causal boxcar filters, and, in the CUE-aligned upper panels, truncated at the center point of the reaction-time range for each coherence. Gray shading denotes standard error of the mean over trials (SEM); not visible in most traces due to large numbers of trials. This neuron shows a clear covariation of firing rate with choice and coherence. Over 1800 trials were used in total for computing the firing rates for this neuron and for each trace more than 100 trials were used for computing the average and the standard errors. **B**: Firing rates of the same neuron shown in A grouped by reaction time. Color now represents average reaction time, from fast (purple) to slow (orange). Other conventions as in A. **C, D:** Firing rates of another neuron in PMd, showing consistent choice selectivity and covariation with coherence (C) and reaction time (D). Conventions for C are the same as A. Conventions for D are the same as in B. This neuron also shows non-monotonicitythat may arise due to the mixing of various signals such as decision-formation and motor preparation associated with a decision-making task.

We performed principal components analysis (PCA) on the set of all firing rate profiles (for both PREF and NONPREF conditions, averaging over strengths of sensory evidence) recorded from the PMd neurons. A schematic for this analysis is shown in Fig. 4A. The first two principal components (PCs) captured ~93% of the variance in trial-averaged firing rates during the decision-making task. That only two PCs capture most of the variance is because differences in evidence, choice, and reaction time are all collapsed in the overall averages; more dimensions are needed when the PCA is based on condition-specific means. However, these PCs were designed to reveal mixed signals that were not necessarily associated with the decision process. Fig. 4B shows the first two principal components estimated in this way from the PMd firing rates. The first PC profile (*X*_1_, ~74% of variance) contributes a monotonic ramp-like change in firing rate in line with the two decision-making models. However, the second PC (*X*_2_, ~19%) rises and then falls, peaking approximately 150ms before the onset of movement.

**Figure 4:**
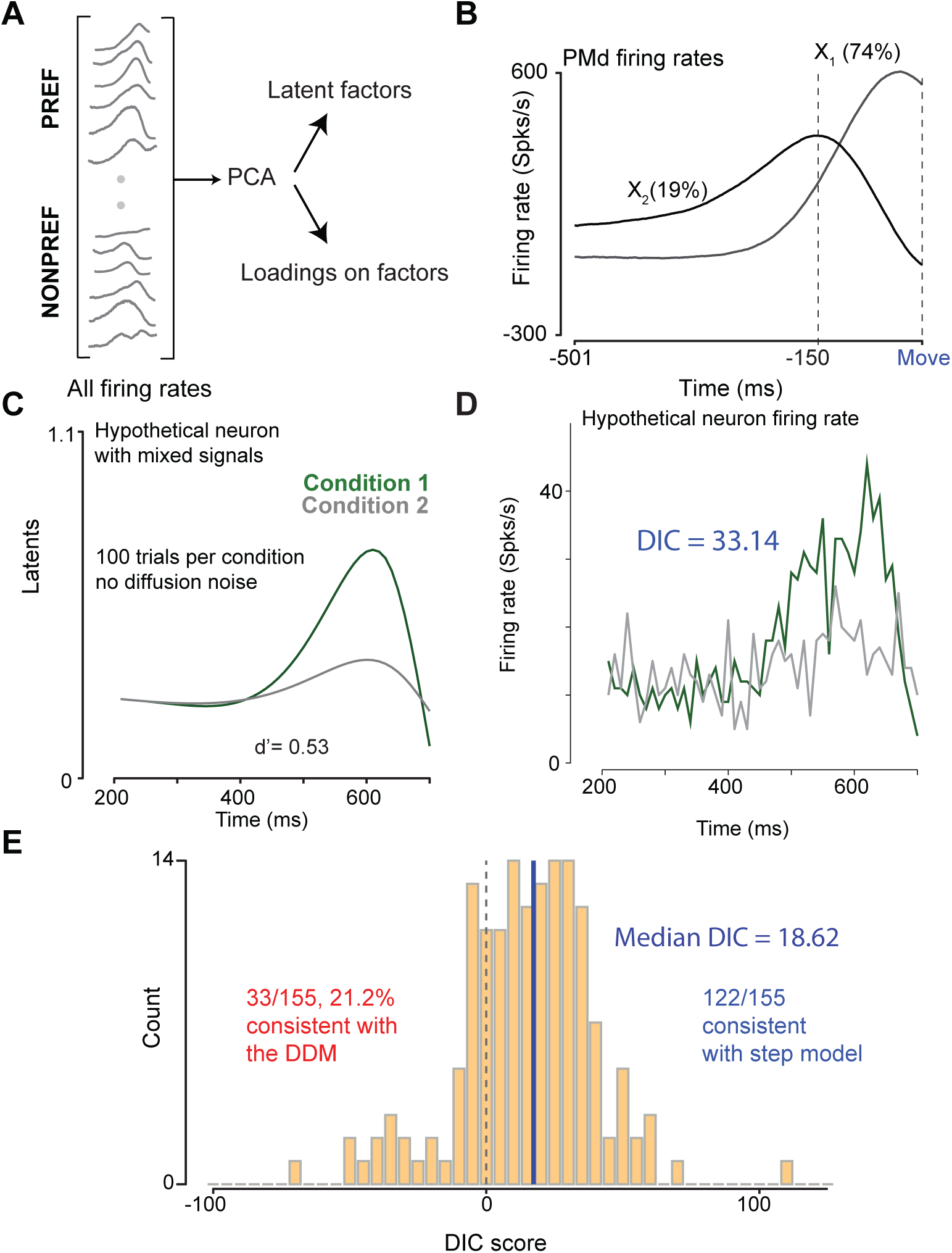
Hypothetical neurons with mixed non-monotonic firing rates are misleadingly classified as stepping. **A**: Schematic of the analysis using principal components to describe the components of trial-averaged firing rates for the PMd neurons during the decision-making task. PCA provides access to the factors (shown in B) and the loadings on the factors. **B:** The first two factors estimated via PCA on the PREF and NONPREF firing rates for PMd neurons. PC1 (*X*_1_) explains ~ 74% of the variance and PC2 (*X*_2_) explains ~ 19% of the variance. *X*_1_ has a monotonic increase in firing rate that is consistent with the DDM and the step models whereas *X*_2_ is essentially inconsistent with both models. **C:** Latent dynamics of a hypothetical neuron meant to mimic the non-monotonic firing rate patterns observed in PMd with components involving both increases and decreases in firing rate resembling the shape of *X*_2_. Two conditions are plotted in green and grey, respectively. For this hypothetical neuron, the latents are identical for every trial and thus plotted on top of one another. Hypothetical arm movements are assumed to occur at t=700 ms. Data are aligned to the stimulus onset. **D**: Trial-averaged firing rates of the hypothetical neuron shown in C. For each trial spike trains were generated using a Poisson model output process with latent dynamics given in C. Firing rates were obtained by averaging over trials for each choice. DIC scores for the neuron shown in C, D supported the step model. Note that no steps were involved in the simulation but the model selection method returns DIC scores that identify the neuron as better described by the step model. **E**: Histogram for the combined DIC score for all 155 neurons simulated with complex firing rates. This distribution also includes neurons simulated with multiple conditions (i.e., multiple drift rates). The distribution is heavy tailed with neurons that arbitrarily get assigned extreme DIC scores both in favor of the DDM and the step model. The median DIC score was in favor of the step model even though no steps were included in the hypothetical neurons.

#### Simulated non-monotonic firing rates are assigned to the step model

We used the profiles identified by PCA as the basis for a simulation (200 trials per condition, 2 stimulus conditions, again one with a strong drift rate and the other with a zero drift rate) with latent firing rate time courses showing increases and decreases broadly mimicking the shape of *X*_2_ (Supp. Fig. 6A). We found that model selection applied to these hypothetical neurons with non-monotonic firing rates tended to favour the step model (Figs. 4C,D). This assignment persisted even when the latents were initially generated from a DDM and only then modified to incorporate time-varying firing rate profiles based on *X*_2_ (Supp. Figs. 6B, C). Of the 155 hypothetical neurons simulated with non-monotonicfiring rate profiles and robust firing modulation, 122 (Fig. 4E, binomial test, 78.71%, p=3.32×10^−13^) were better described by the step model even though the simulation process introduced no explicit steps.

#### Non-monotonicity in PMd data predicts DIC score

Returning to the data, we asked if DIC score was related to the degree of non-monotonicity in PMd firing rates. We binned neurons by their loadings on *X*_1_ and *X*_2_ averaged over both PREF and NONPREF trials, and estimated the percentage difference in assignments to the DDM and step model for each bin in this two dimensional grid (Fig. 5). When loadings were positive on *X*_1_ and negative on *X*_2_ (dashed red ellipse in Fig. 5), firing rates of single neurons were both closer to monotonic and more likely to be selected as consistent with the DDM (Examples 1–3 in Fig. 5). Conversely, for neurons with loadings positive on *X*_2_ and negative on *X*_1_ (dashed blue ellipse), firing rates appeared non-monotonic and DIC scores were more consistent with the step model (Examples 4–6 in Fig. 5).

**Figure 5:**
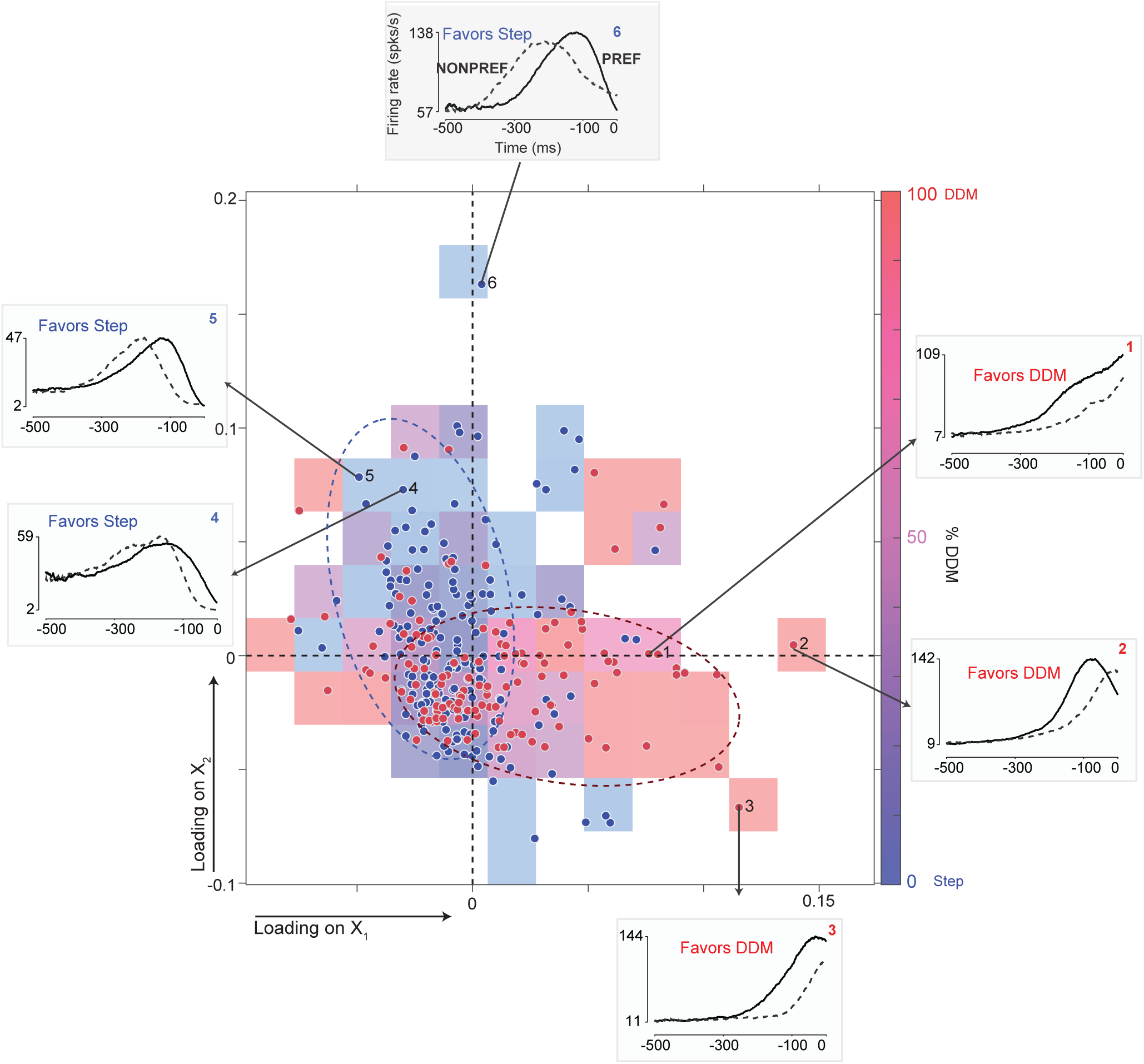
Non-monotonic firing rates in PMd are often labeled as stepping. Classification of neurons as ramping or stepping as a function of firing rate loadings on principal components 1 (*X*_1_) and 2 (*X*_2_). Each colored square represents neurons that fall within a single bin in the two-dimensional loading space. The hue of the square represents the proportion of these neurons classified as ramping from blue (none) to red (all). The saturation varies with the number of units in the bin. Empty bins are shown in white. Blue and red dashed ellipses are drawn to indicate regions of consistency with the step model and the DDM respectively. Dots represent individual neurons colored red or blue depending on whether they are consistent with the DDM or the step model. Insets show firing rates of example neurons for PREF (solid) and NONPREF (dashed) trials, leading up to movement onset at 0 ms. Higher loadings on *X*_1_, which is broadly consistent with a monotonic increase in firing rate, lead to greater consistency with the DDM (red ellipse and insets 1–3). Higher loadings on the non-monotonic *X*_2_ are associated with more frequent assignments to the step model (blue ellipse and insets 4–6).

Consistent with this qualitative picture, the average loading for a neuron on *X*_2_, again averaging over PREF and NONPREF trials, was positively correlated with the DIC score. Thus greater non-monotonicity, as modeled by projection onto *X*_2_, is associated with a greater likelihood of being assigned to the step model (Spearman’s *ρ* = 0.16, p=.006).

We also found a negative correlation with the average loading on the first PC (*X*_1_) suggesting that neurons with stronger overall changes in firing rate are likely to be better described by the DDM (PREF: Spearman’s *ρ* = −0.18, p =0.0015). These correlation analyses were also consistent with a non-parametric regression that attempted to predict DIC score as a function of the loadings on *X*_1_ and *X*_2_ (Birkes and Dodge, 2011, F(308) = 19.85, p < .00001, 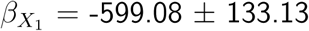, t(309)=−4.99, p < .0001; 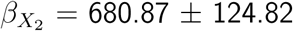, t(309) = 5.46, p < .0001).

Recall that neither the DDM nor the step model predicts non-monotonic firing rates profiles, and so this additional signal mixed in the PMd data should not overtly favour either model during selection. Nonetheless, we find that it has a clear impact on outcomes.

Together, the analyses suggest that non-monotonicity in firing rate profiles can impact the results of model selection. Neurons with more non-monotonicity in their firing rates were more likely to be identified by model selection as more consistent with the step model.

### 3. Brittleness to parameter non-stationarity or adaptation

Model selection, like other statistical analyses applied to high-dimensional complex systems, requires a great deal of data to yield confident results. This requirement increases the chances that a system will adapt or experience fluctuations over the period spanned by data collection, or that trials collected under different experimental conditions will need to be grouped together. In this case, even when a system is well described by a single parametric *form* of model, the specific parameter *values* that best characterise it may vary with time or experimental factors. In our final study we asked whether model selection would prove to be robust to such variation when the models tested formally assumed a single setting of the parameters to explain all data, albeit one that is treated as unknown and modelled with uncertainty in a Bayesian analysis.

Parameter variation might arise naturally within a neural implementation of a drift-diffusion process where evidence for opposite decisions is accumulated in two competing neural populations (e.g., Usher and McClelland, 2001). In such circuits, evidence supporting a NONPREF decision arrives indirectly through inhibition from the competitor neural pool. If the gain of such inhibition differs from that of the direct positive evidence then both the drift rates and the diffusion variance for NONPREF trials would be different from those in the PREF condition. Indeed, it may even be that ongoing inhibition is weak or non-existent, so that the two populations are engaged in a ‘race’ to separate thresholds, with a strong inhibitory signal unleashed by the population that wins. In this case, individual neurons might appear to “step” down for NONPREF decisions, but still “ramp” up in PREF conditions. Differences in dynamics between the PREF and NONPREF conditions have been suggested previously for both PMd (See Fig. 6 of Thura and Cisek, 2014) and LIP (Fig. 7 of Roitman and Shadlen, 2002). We considered these two possibilities in two simulations (Fig. 6 and Fig. 7).

**Figure 6:**
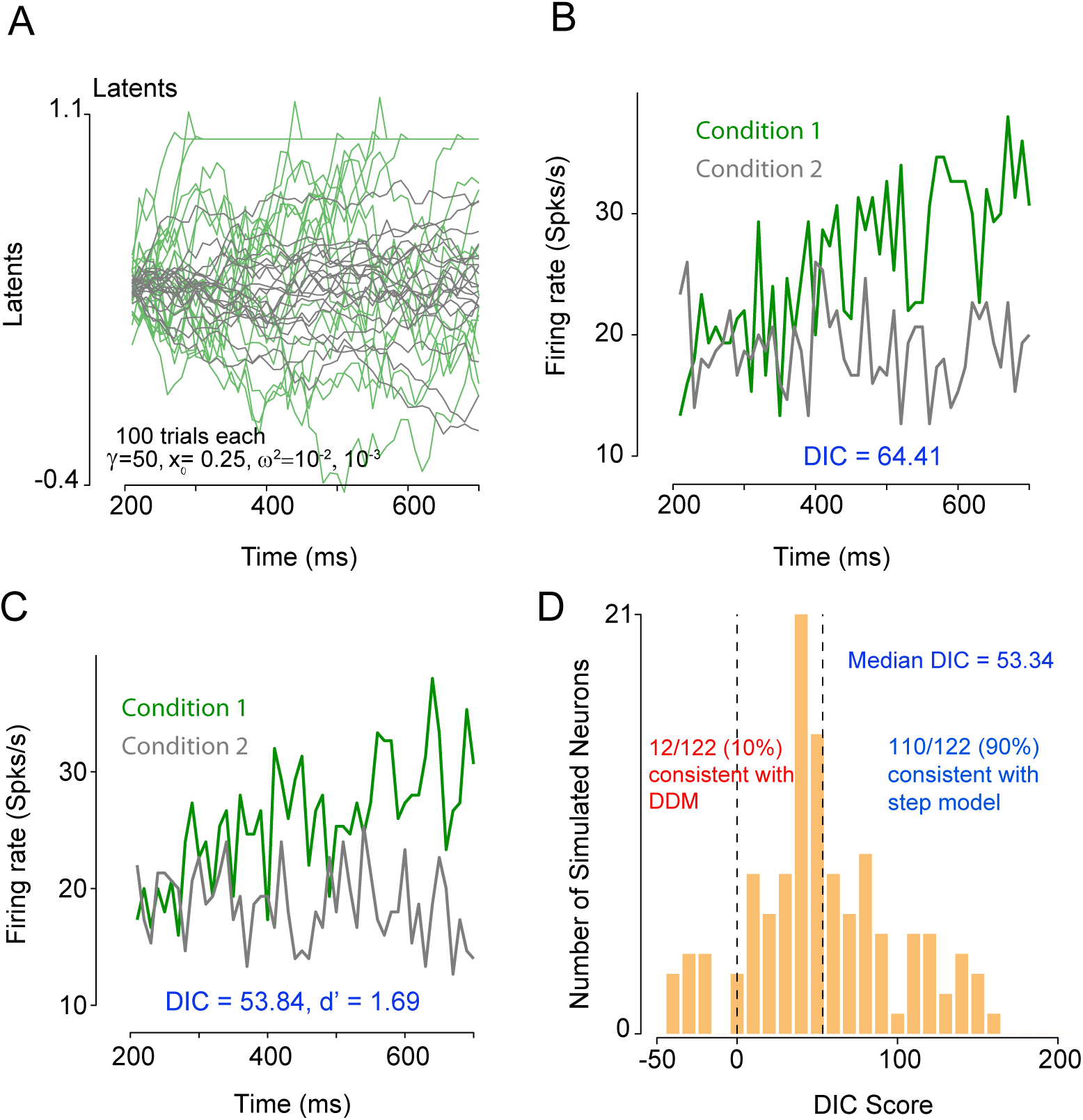
Hypothetical neurons with non-stationary diffusion noise are classified as stepping. **A:** Dynamics of latent variable (including absorbing upper bound) for a hypothetical neuron with DDM dynamics and condition-dependent diffusion variance. Condition 1 (green traces) had *ω*^2^ = 10^−2^; condition 2 (gray traces) had *ω*^2^ = 10^−3^. **B**: Firing rate for the simulated neuron shown in A. DIC score for this hypothetical neuron was consistent with the step model even though no steps were introduced. **C:** Firing rates of another example simulated DDM-based neuron with condition-dependent diffusion variance as in A and B, classified as stepping. **D**: For the majority of the neurons simulated from the DDM with condition-dependent diffusion variance DIC scores favoured the step model. Variance for condition 1 was set at 10^−2^; for condition 2 it was either 10^−3^ or 10^−4^.

#### Difference in diffusion variance for different conditions

In the first case, we simulated responses in which one set of trials had a high drift rate and diffusion variance on the same order of magnitude as that estimated for the LIP data, while the other had zero drift and diffusion variance was either one or two orders of magnitude smaller than the first. This setup was broadly consistent with the case of inhibition arriving with low gain from a competing population, although in our simulations the gain depended on the overall signal rather than the instantaneous input. Two example simulations where the diffusion variance for one set of trials was one order of magnitude smaller than the variance of another set of trials are shown in Figs. 6A-C. These hypothetical neurons with latent dynamics from the DDM but different values of diffusion noise were often identified as being more consistent with the “step” model, even though no steps were present in the generative model (Fig. 6D, 110/122 simulated neurons with were identified as consistent with the step model, p=2.73×10^−21^, binomial test).

#### Drift-diffusion for PREF choices, and steps for NONPREF choices

The second simulation was analogous to the ‘race’ model of decision formation described above, where the neurons would appear to ramp for PREF choice trials but step down on NONPREF choice trials. Figs. 7A, B and Fig. 7C show examples of two hypothetical neurons simulated using this hybrid model. As the numbers of PREF and NONPREF trials in the simulations are equal, robust model selection would arguably generate scores distributed near to and around 0, with roughly half of the hypothetical neurons being assigned to each class. Alternatively, given the robust modulation modeled for the PREF choices and relatively weaker modulation (and thus less “signal”) for NONPREF choices, it might also have been reasonable for DIC scores to modestly favour the DDM. In fact, model selection assigned *all* of the simulated hypothetical neurons to the step model (Fig. 7D, median DIC score=197.71), even when the DDM parameters for drift and diffusion were substantial in the PREF direction.

**Figure 7:**
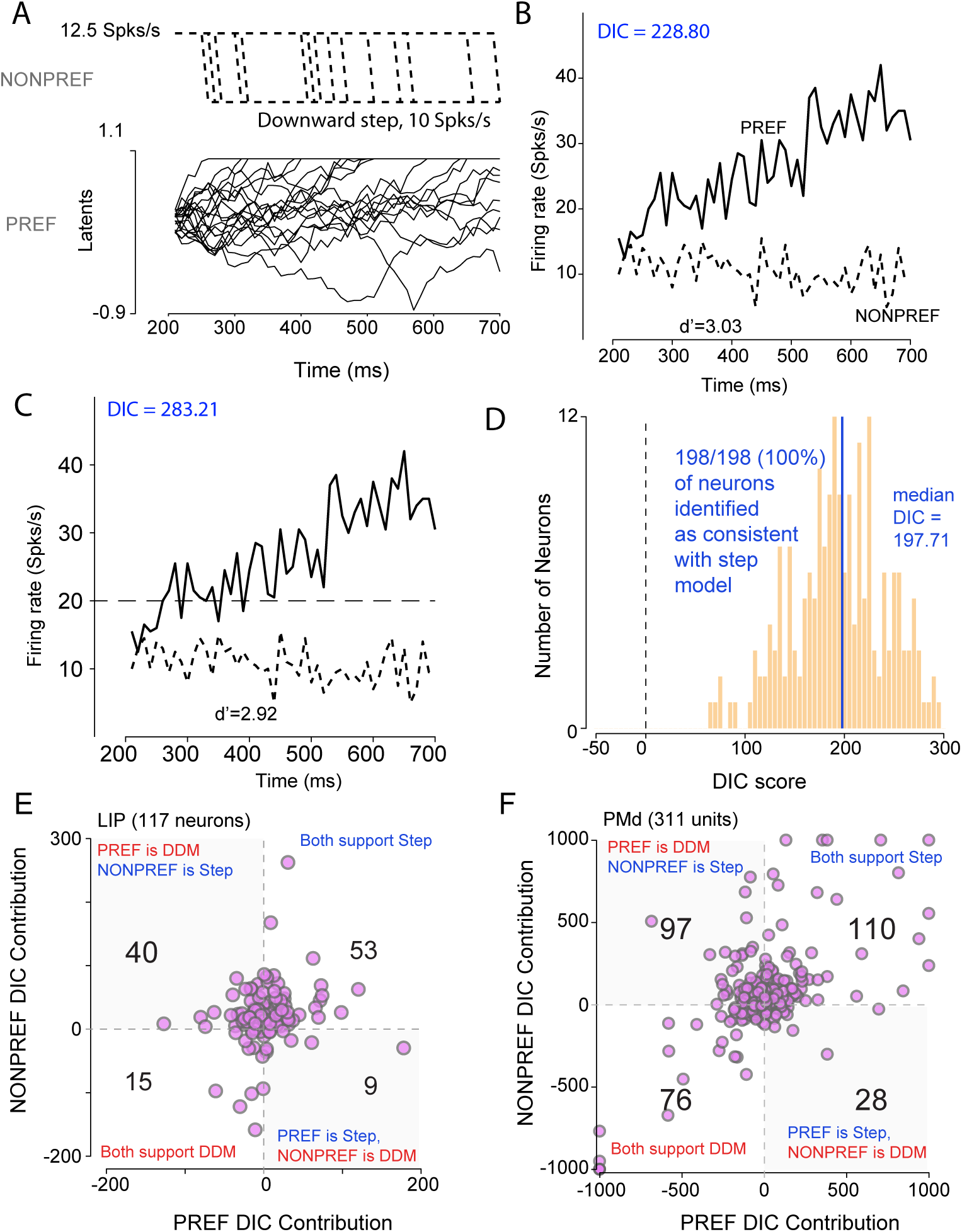
Hypothetical neurons with different models for different choices are identified as stepping. **A:** Latent dynamics of a hypothetical neuron with PREF direction latents described by a DDM and NONPREF direction latents described by the step model. **B**: Trial-averaged firing rates of the hypothetical neuron shown in A. PREF choice trials are shown in solid lines and NONPREF choice trial are shown in dashed lines. For each trial spike trains were generated using a Poisson model output process with latent dynamics given in A for the two different choices. Firing rates were obtained by averaging over trials for each choice. DIC scores for the neuron supported the step model. **C** Firing rates of another hypothetical neuron with DDM latent dynamics for PREF choices and step for NONPREF choices but identified by model selection as consistent with the step model. Conventions as in B. **D**: Histogram for the combined DIC score for all 198 neurons simulated with DDM for the PREF direction and the step model for the NONPREF direction. The median DIC score overwhelmingly favored the step model even though the units had large PREF direction decision-related responses that involved ramping. **E**: Plot of contributions to the DIC score from PREF trials and NONPREF trials for each pseudo-independent neuron in the LIP dataset. For 49/117 neurons, the DIC scores from the PREF and NONPREF trials did not agree. The number of neurons consistent with each combination of PREF and NONPREF DIC contribution is provided as text in each quadrant. **F**: Same as E except for PMd data. 125/311 units had DIC score contributions that were different between PREF and NONPREF trials. Again, many PMd units have DIC score contributions more consistent with the DDM for PREF compared to NONPREF conditions. DIC score contributions outside ± 1000 are clamped at ± 1000 to highlight structure for other units.

PREF and NONPREF trials combine to affect DIC scores in two ways. First, the parameters of the competing models are fit to all trials at once, and so where the two groups are generated by different processes the resulting posterior distributions must reflect a compromise between conflicting sources of evidence. Second, the final DIC score is formed by summing contributions from each trial to the deviance and its posterior average (see Methods).

If both PREF and NONPREF trials are generated from a single model with a single set of parameters, then we expect the net contributions to DIC from both groups of trials to be consistent to within noise-driven variability. We wondered if the converse would be consistently true. That is, in these neurons simulated from this mixture of the DDM and the step model, would the DIC contributions made by the corresponding groups of trials correctly identify that the PREF trials come from the DDM and NONPREF trials come from the step model (even though the parameters would still be fit to all the trials together). This was indeed the case for a subset (32/198) of the hybrid model simulations, with the sum of PREF trial contributions pointing to the DDM model, but NONPREF contributions supporting the step model. However, for the majority of the hypothetical neurons (166/198), the DIC score contributions from both the PREF and NONPREF directions favoured the step model. Thus, although systematic inconsistency in the direction of the contribution made by two subsets of the data (here PREF and NONPREF trials) may be reasonably taken to challenge the DIC results, the converse does not hold. Consistency in the directions of net contributions from two groups of trials does not imply a robust conclusion.

#### Results of model selection on PMd and LIP data differ for PREF and NONPREF trials

We wondered whether the neural data might also show signs of inconsistency between PREF and NONPREF trials. We first examined contributions to the total DIC score made by each group (Fig. 7E, F). If each neuron was equally well fit by ramp or by step model in both groups of trials then we might have expected these contributions to have been well correlated. Instead, we found a broad diversity of patterns, with 49/117 (41.88%, Fig. 7E) neurons in LIP and 125/311 (40.16%, Fig. 7F) units in PMd exhibiting inconsistency in the contributions from PREF and NONPREF trials. In general, PREF trial contributions pointed to the DDM more often than those of NONPREF trials for both LIP (Chi-Squared test, PREF: 55/117 support DDM, NONPREF: 24/117 support DDM, *χ*^2^(1) = 19.06, *p* < .0001, Fig. 7E) and PMd (Chi-squared test, PREF: 173/311 support DDM, NONPREF: 104/311 support DDM, *χ*^2^(1) = 30.92, *p* < .0001, Fig. 7F).

## Discussion

We asked whether model selection could robustly distinguish between two conceptual hypothesis classes given the inevitable heterogeneity of data from complex systems. In our simulations—and those of Latimer et al. (2015b)—when data were generated from an exact instance of one or the other of the two mathematical models being compared, the DIC criterion correctly favoured the generating model. In this way, then, model selection worked as intended. However, once the parameters of the generating process departed from the formally specified DDM-based model chosen for analysis, model selection proved brittle and mis-assignments became common. This happened even for intuitively minor departures that preserved the gradual changes in single-trial firing rate that are characteristic of the “ramp” class that the DDM was meant to represent, without introducing the abrupt changes key to the competing “step” model. The neural data from decision-making experiments that we examined shared diagnostic features with the variant models we simulated, raising concerns about the interpretability of model selection results in these data.

Information criteria for model selection that approximate a probabilistic distance between models and data are intended to be robust to small variations in the generative process. When neither model fits exactly, the information-theoretically closer of the two models to the data should also be the more conceptually consistent with them. Our results belie this common assumption, demonstrating empirically that model selection applied to complex data can be brittle. Although we only discussed results with DIC here so as to remain comparable to earlier work, we found much the same to be true of the “widely applicable” information criterion (Watanabe, 2010). Rather than depending on the precise criterion chosen, it seems likely that brittleness arises from a fundamental mismatch between probabilistic or information-theoretic proximity and conceptual consistency.

The situation can be summarised using a fruit analogy (Fig. 8). Suppose that we seek to classify a sample fruit into either the pome or the citrus families (model classes) based on its appearance alone. Both families are difficult to describe in their respective entireties: the potential range of hybrid or sport pome and citrus fruit is so extensive and complex that no simple model of their appearance will capture every potential variation. Instead, we are forced to choose two representative instances against which to compare the new fruit. Call them the apple and the orange (Fig. 8A). When presented with a new fruit we ask whether it appears closer to our model of an apple than an orange, and if so classify it as “pome”. This is the operational essence of model selection as used in the complex sciences. For instance, in the case we considered here, the pomes are steps, the apple the Poisson change-point model (with all its constraints), while the citrus are ramps and the orange the drift-diffusion-based model.

**Figure 8:**
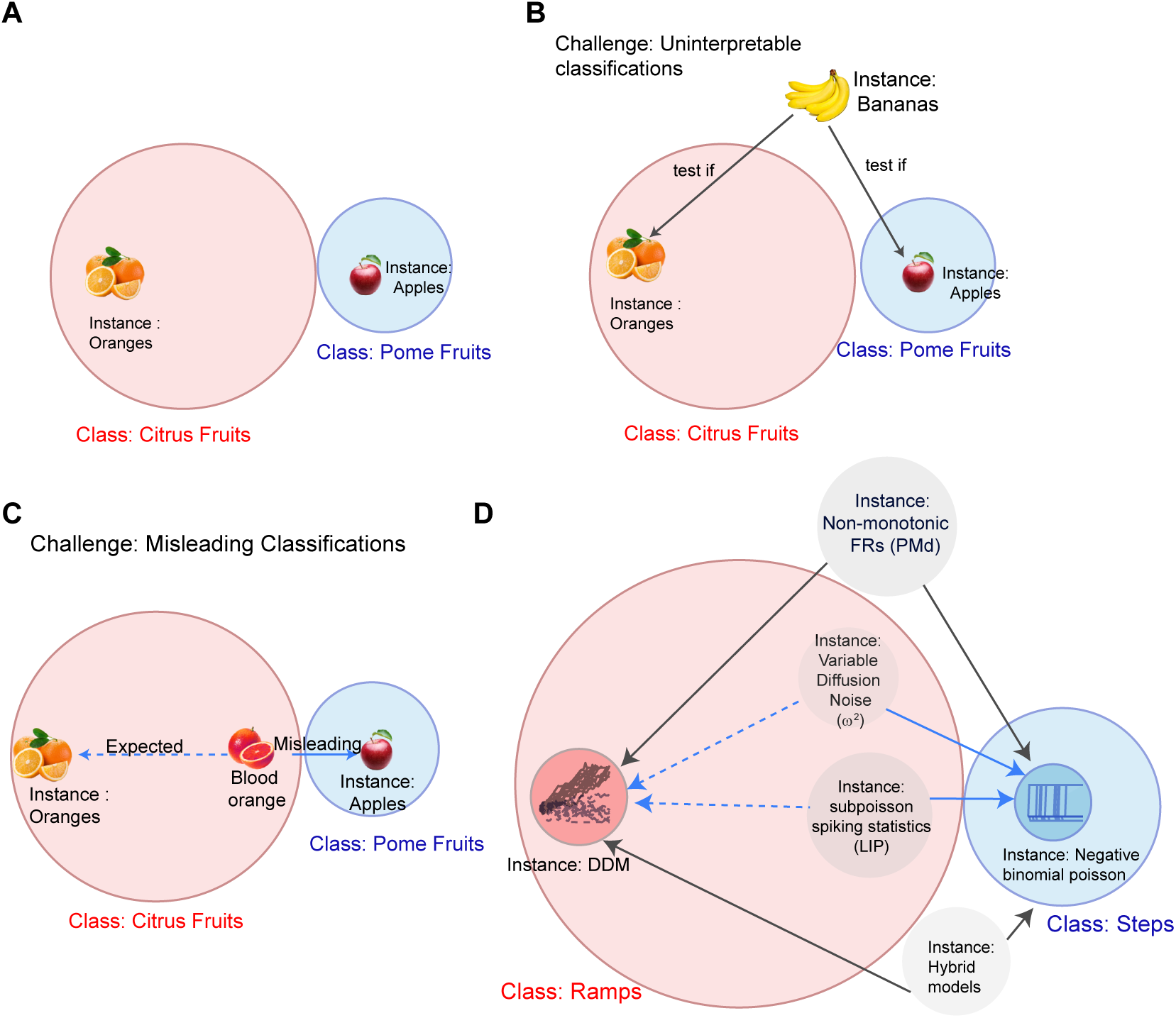
A fruit analogy summarizing the challenges in model selection. **A:** A cartoon summary of the use of model selection applied to complex natural systems. Conceptual model classes must be exemplified by a mathematical model. In the analogy, the model classes of citrus and pome fruits are represented by oranges and apples respectively – each only a single instance of the wide range spanned by the corresponding family. **B:** Model selection applied to data (here a banana) that lie outside both model classes can produce uninterpretable results. Answers to the question “Is a banana more like an apple or an orange” are unlikely to be helpful. **C:** Misleading results can arise when legitimate members of one model class (here a blood orange) happen to appear closer to the exemplar of the other one (the apple). Such brittleness depends partly on the metric chosen (here, one that weights similarity of color over form) and partly on the narrowness of defined exemplars (here, an orange class that allows little variation in color). However, these issues are systemic to model selection. **D:** Our simulations examined the results of model selection assigning neural firing dynamics to the conceptual classes of “ramps” and “steps”, using the exemplars of a particular drift-diffusion model and negative-binomial step-time model respectively. Simulated neurons with mixed responses imposed over an otherwise DDM-consistent generative process were consistently assigned to the step model. A similar outcome emerged for neurons generated from a hybrid model with steps for one set of trials and the DDM for another set of trials. Other simulations derived from the DDM were more clearly within the “ramps” class: modifying only the form of spiking noise, or allowing condition-dependent diffusion variance. These were misleadingly assigned to the “steps” class.

One challenge posed by this model selection approach is obvious. Our procedure identifies every fruit as “pome” or “citrus”. If applied to, say, a banana, such a classification is meaningless; but the model selection procedure has no automatic way of rejecting the comparison (Fig. 8B).

More pernicious, perhaps, is the case of a sanguinello or blood orange. Intuitively and botanically this is a citrus fruit. However, if the model of “orange” appearance is specified to have a very narrow range of possible skin colours, while that of “apple” tolerates many hues (green, yellow, red,…), then it is possible that when forced to select between these two specific models the “apple” is found to be closer (Fig. 8C). Arguably, this is analogous to what happened in many of our case studies (Fig. 8D). Data generated from versions of the DDM that retained the gradual firing rate change over the trial that is central to the informal definition of “ramp” nonetheless fell outside the narrow definition of the model instance used for testing, and so were classified as a “step”.

### Consequences for interpreting decision-making processes

We were led to consider the brittleness of model selection while attempting to extend the step or ramp characterisation of decision-related activity discussed by Latimer et al. (2015b). However, as our results demonstrate, model selection in this application proves to be brittle in many respects. Although we believe the underlying question to be well-founded and important, the particular model instantiations for step and ramp chosen in that study fail to represent the intended conceptual model classes fully. In particular, apparently minor deviations from the ramp DDM model lead to mis-assignments of model selection, and evidence for such deviations is evident in neural data. Based on these concerns, we cannot rule out the possibility that, despite the apparent model-selection evidence from Latimer et al. (2015b), all neurons in LIP (and PMd) with decision-related activity reflect some form of ramp-like drift-diffusion process.

### Recommendations to mitigate the brittleness of model selection

Despite the challenges to its application and interpretation, we believe that model selection should still play an important role in systems neuroscience (Churchland and Kiani, 2016; Durstewitz et al., 2016; Linderman and Gershman, 2017). To paraphrase Box’s position: all models of a complex system are inescapably wrong, and so hypothesis testing will always lead to rejection in the face of enough data. Nonetheless, scientific progress depends on identifying models that are both reasonable and useful.

An increasingly sophisticated arsenal of data-gathering technologies (Jun et al., 2017; Maynard et al., 1997), along with novel capabilities to intervene with ever finer precision in the central nervous system (Lee et al., 2015; Packer et al., 2014), make it possible to test a widening array of models by evaluating qualitative, rather than quantitative, predictions that are able to distinguish between entire classes of conceptual model. Where they are possible, such experiments may obviate the need for formal model selection. But for many models, including those represented by the “step” and “ramp” considered here, such qualitative approaches remain technically intractable.

How might a future model-selection study then mitigate these challenges? Based on the insights from the case studies here, we offer three recommendations that could help make conclusions based on model selection applied to neural responses more robust.

First, an initial set of qualitative methods should be used to rule out any potential instances in the data that are broadly inconsistent with all of the working hypotheses being considered—that is, “bananas” in our fruit analogy. This may require a close look both at the overall behaviour, and at the variations within the data set. Where the data are single neuron recordings, for example, it may be that a simple summary statistic such as the grand-mean firing rate over all trials and neurons appears broadly consistent with the models under consideration, but that the contribution of individual neurons to this mean are not. Although apparent departures from the hypotheses in any one neuron might be put down to noise, systematic deviations that appear in some degree in many neurons suggest that the model space needs enlargement. Such systematic deviations may be revealed by methods such as PCA that examine the pattern of firing rate profiles of neurons in the population as a whole. This was the case in the PMd data set considered here. The grand-mean firing rate profile across neurons appeared to increase gradually throughout the trial (Supp. Fig. 2) in a manner plausibly consistent with either step model or DDM. However, PCA revealed that this grand mean concealed a significant component of variation in individual neuron firing rate patterns that did not fit either hypothesis; and we found that the strength of this component in an individual neuron firing rate was predictive of the model selection outcome.

A second proposal is to carry out confirmatory qualitative analyses after selection, to ask whether the quantitative assignments appear plausible—and so to test for the possibility that a blood orange has been classed as an apple. This might involve reassessing properties of the data themselves in light of the selected model fit, although we did not take such an approach here. Instead, we observed a qualitative feature in the model selection results themselves which cast doubt on robustness. If model selection were robust, then—for data that are collected in trials, as ours were—it would be reasonable to expect that each type of trial should influence the selection result in the same direction, even if not with the same confidence. In fact, for many of our PMd and LIP neurons PREF and NONPREF trials systematically disagreed in their contributions to selection, with the overall assignment depending on which contribution was the stronger. In principle, this finding could mean that the different groups of trials were, in fact, better described by different classes of model. However, we found that a similar situation arose in simulations when all the trials were generated by a single class of model but with different parameters. Thus, the observed heterogeneity suggests again that the model space chosen is too restrictive and so the results of selection suspect.

Both recommendations above are diagnostic of brittleness, but do not in themselves offer a clear solution. In fact, however, both point to the conclusion. Despite the appeal inherent in selecting a single, elegant mathematical model to represent each working hypothesis under consideration, model selection results will align more closely to the intuitive hypothesis classes if each is represented by a composite model which attempts as far as possible to tile the range of the hypothesis space — representing “citrus” by oranges, lemons, limes, grapefruit and more exotic varieties; and “pome” by apples, pears, quince, loquat and others (Burnham et al., 2011). In the current example, for the “step” class one might consider models that allow more than one transition between the parametrized firings, and trial-to-trial fluctuations in gain (Goris et al., 2014); while “ramp” models, even if restricted to the DDM, may incorporate variable start times for different trials (Kiani et al., 2014), a non-zero lower bounding rate (Zylberberg and Shadlen, 2016), and elements such as urgency (Ditterich, 2006) or bound collapse (Hawkins et al., 2015b,a). Both models may be elaborated with different spiking statistics (Pillow, 2009).

None of these proposals provides a fully automated fix to the issues we have raised. However many qualitative measures have been evaluated, it is always possible that another measure will reveal the assigned fruit to be an imposter. Any enumeration of formal models within a hypothesis class — say of all the varieties of tangerine — will always be open to the discovery or invention of another. Thus, formal statistical model selection cannot be more than one tool of many, that must be combined to advance our understanding of complex systems such as the brain.

## Summary

Fields such as systems neuroscience, which study heterogeneous and complex natural systems, face a particular challenge in the evaluation of theoretical models. All such models will inescapably be rejected by hypothesis testing based on enough data. So, scientific progress depends on identifying models that are reasonable, and more importantly useful, rather than “correct”. In neuroscience, improvements in data gathering and interventional technologies increasingly make it possible to test qualitative predictions of some models. But such methods still have limits, and there remains a need for formal model selection where such tests are intractable. While we broadly support the use of such methods, we have identified a key challenge which may lead to misguided conclusions about the neural dynamics underlying behavior. To guard against such challenges, model selection should employ a plurality of model instances within each broad class to be considered, and should be combined with qualitative measures of model agreement to provide a more robust path to scientific progress.

## STAR Methods

The task, training and electrophysiological methods used to collect the data used here have been described previously (LIP: Rorie et al. 2010; PMd: Chandrasekaran et al. 2017) and are reviewed briefly below. The methods for fitting the models are elegantly and comprehensively described in the recent report (Latimer et al., 2015b) and book chapter (Latimer et al., 2015a) that developed the model selection framework. Again, we review these only briefly.

### Oculomotor visual discrimination task and recordings from LIP

#### Task

Two monkeys (Ar and Te) performed a variant of a *fixed duration* random dot discrimination task where both stimulus difficulty—set by the coherence level of the random dots—and reward contingencies were manipulated (Supp. Fig. 1A, Rorie et al., 2010). The monkeys were trained to detect the net direction of motion of a noisy moving random-dot stimulus and report it by making a saccadic eye movement to one of two targets positioned in line with the axis of motion being discriminated. The dots stimulus was displayed for 500 ms. Following this, monkeys were required to maintain fixation for a variable delay (300–550 ms), and were then cued to initiate the eye-movement report. A correct response was rewarded with a drop of juice; the size of the reward depended on the value assigned to that target.

#### Recordings

While monkeys performed this task, single neuron activity was recorded from LIP using single, sharp dura-piercing electrodes. The original report included 81 LIP neurons (51 from Monkey Ar, 30 from Monkey Te) and neurons were selected in the original study using a variant of the delayed saccade task. Each neuron was tested in 4 different reward conditions. We separated these conditions, giving a pool of 324 pseudo-independent recordings (81 neurons × 4 reward contingencies) with which to examine the model selection technique.

#### Model selection analysis

For analysis of these LIP firing rates (FRs) we took all data in an interval from 200 ms after dots onset until 200 ms after the dots offset; the lag was designed to avoid the initial transient often seen in LIP responses and to approximate the latency with which visual information arrives in LIP (Roitman and Shadlen, 2002). As in Latimer et al. (2015b), we only selected firing rates from recordings with adequate choice selectivity (d′ > 0.5, LL: 36, HH: 26, LH: 33, HL: 22). This selection criterion left a total of 117 pseudo-independent recordings. Following Latimer et al. (2015b), we binned trials in the LIP data according to their signed motion coherence into six different groups by level (3 in each direction), with 0% coherence trials placed in a separate group.

### Somatomotor reaction time visual discrimination task and recordings from PMd

#### Task

Two trained monkeys (Ti and Ol) performed a visual reaction time discrimination task (Chandrasekaran et al., 2017). The monkeys were trained to discriminate the dominant color in a central static checkerboard composed of red and green squares and report their decision with an arm movement (Supp. Figs. 1B-D). If the monkey correctly reached to and touched the target that matched the dominant color in the checkerboard, they were rewarded with a drop of juice. The reaction time task allowed monkeys to initiate their action as soon as they felt they had sufficient evidence to make a decision. On a trial-by-trial basis, we varied the color coherence of the checkerboard, defined as 100 × (*R* − *G*)/(*R* + *G*). (Supp. Fig. 1D), where R is the number of red squares and G the number of green squares. The color coherence value for each trial was chosen uniformly at random from 14 different values arranged symmetrically from 90% red to 90% green. Reach targets were located to each side of the checkerboard. In many sessions their colors were also chosen randomly on each trial; in a few other sessions the target configuration was fixed over a block of contiguous trials.

#### Recordings

In our original study, we reported the activity of 996 units recorded from Ti (n=546), and Ol (n=450) while they performed the task (Chandrasekaran et al., 2017). PMd units demonstrate temporal complexity in their firing rate profiles. We analyzed an 806 unit subsample and applied the visuomotor index of Chandrasekaran et al. (2017) that measures the degree of sustained activity to separate this population into the broad categories of increased (429/806), perimovement (118/806) and decreased (259/806) units. **Increased units** exhibited ramp-like increases in average firing rate, time-locked to the onset of the visual stimulus, with slopes that varied with stimulus coherence. These are the classic candidates for neurons that might carry an integrated decision variable for the discrimination task (Supp. Figs. 2A,B). We therefore focused for this study on these increased units, again filtering for significant choice selectivity (d′ > 0.5), arriving at 311 PMd units for analysis.

#### Model selection analysis

For analysis of single-trial firing rate dynamics, we used spike trains beginning 100ms after the checkerboard onset and extending until the initiation of movement. The 100 ms assumed latency matched that found in population analysis in our previous study. Analysed spike trains were therefore approximately 200 to 900 ms long, and the probability density of the time analysed for each trial roughly matched a gamma distribution. To simplify and speed up the model selection analysis, we first created a signed directional evidence measure signal which combined the checkerboard signed coherence and the target configuration. We then grouped the 14 signed directional evidence values into six different levels. {−90, −60, −40}, {−30, −20}, {−10, −4}, {4 10}, {20,30}, {40, 60, 90}.

### Statistical Analysis

#### Regression analysis for LIP neurons

We pursued two regression analyses to examine the relationship between under dispersion and DIC scores for the LIP neurons.

In the first, we tested the relationship between DIC and Fano factor for the NONPREF choices (*FF_NONPREF_*).

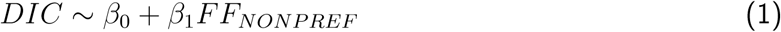

In a second regression analysis, we included both the starting and the ending firing rates for the PREF and NONPREF choices.

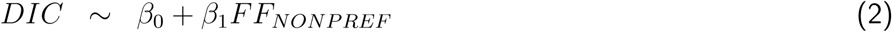

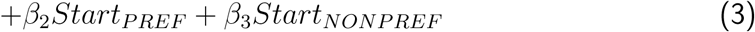

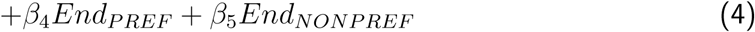

We used standard regression methods to fit these models and report the model fit, the coefficients, t-statistics, and p-values in the main text. We also report the partial *R*^2^ for the regression between DIC and NONPREF Fano factor, after accounting for the starting and ending firing rates. Using other intervals (e.g., the entirety of the dots epoch) for the Fano factor did not materially alter the results nor did using the PREF direction Fano factor in place of the NONPREF Fano factor.

### Principal component analysis of PMd neurons

To perform principal component analyses (PCA) of the firing rates of PMd neurons, we aligned the data to movement onset and averaged to obtain the mean PREF and NONPREF response profile for each one. We then created a large matrix of firing rates of size 2*N* × *T*, where *N* refers to the number of units analyzed and *T* the time period. We chose the 500 ms before movement onset as the time period for the PCA analysis.

We first subtracted the mean across neurons from this matrix and computed the principal components via eigenvalue decomposition of the covariance matrix of this centered firing rate matrix. We then used the loadings on the principal components for our analyses.

### Correlation and regression analyses for PMd neurons

We used two methods to investigate the relationship between the loading on principal components and DIC score. For the PMd data, we obtained significant numbers of very large outlier DIC scores (> ±1000) either in support of the DDM or the step model. These outliers can impact the values of Pearson correlation and linear regression coefficients. We therefore used Spearman rank correlations and non-parametric regression to investigate relationship between DIC and the loadings (Birkes and Dodge, 2011). For correlation analyses, we used Spearman’s correlation between the average loading on a principal component for the PREF and NONPREF trials and the DIC score for a neuron. We also used a non-parametric regression where we predicted DIC score using the average loading for PREF and NONPREF firing rates on *X*_1_ and *X*_2_ and report the F statistic for the regression as well as t statistics for the predictors (Birkes and Dodge, 2011).

### Model specification for analysis of binned single neuron responses

We briefly review the formulation and the parameters for the step and DDM models introduced by Latimer et al. (2015b,a). The model selection method evaluates whether single-trial responses of decision-related neurons are better explained by a step model or a DDM. Both models assume that an observed spike train is a Poisson process with a rate governed by a noisy, unobserved latent process. Therefore, both the DDM and the step model are “doubly stochastic”. In the case of the DDM, the latents follow a noisy diffusive process whereas in the step model the stochasticity arises from randomness in step times across trials.

#### DDM

The drift-diffusion model of single-trial firing rate dynamics is parameterized as follows. The time-varying firing rate *r_j_*_,1_…*r_j_*,*T_j_* for trial *j* of length *T_j_* time steps (each of size Δ*t*) is determined by a latent trajectory (*x_j_*_,1_…*x_j_*,*T_j_*), which is distributed according to a discrete-time drift-diffusion process. The latent process starts at an initial state *x*_0_. At each time step, it evolves with a drift rate of *β_c_*_(_*_j_*_)_ where *c*(*j*) indexes the coherence on trial *j*, and diffusion noise of variance *ω*^2^. The firing rate *r_j,t_* follows this drift-diffusion process until it reaches an absorbing upper bound, given by *γ*. There is no absorbing lower bound. The model for trial *j* can be written as follows (c.f. Latimer et al., 2015b):

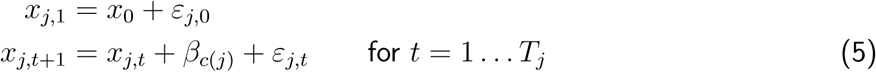

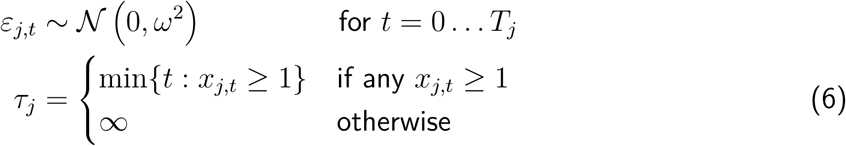

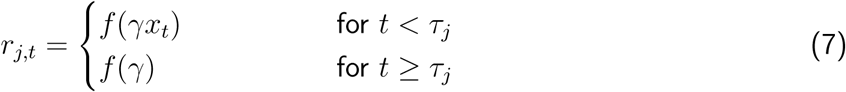

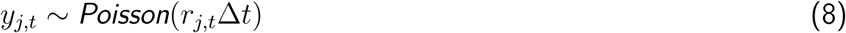

*τ_j_* is the bound-hitting time (the first time bin at which *x_j,t_* ≥ 1). The latent state is converted into a firing rate using the soft-rectification function:

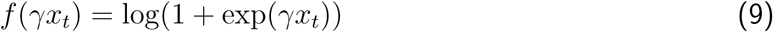

In this formulation, the effective absorbing bound for the latent variable *x* is 1, and the latent is scaled by *γ* to obtain the neuron’s spike rate. This spike rate is then used to generate spike counts from a Poisson distribution. (Note that latent trajectories in Fig. 2, Fig. 6 and Fig. 7 are shown with the bound applied directly to *x*.) The parameters for the diffusion model are: Θ = {*x*_0_*, ω*^2^*, γ, β*_1_…*β_C_*}. The condition-specific slope parameters *β*_1_…*β_C_* allow the rate of accumulation to vary with the strength of the sensory evidence–in our task, given by the coherence level of the random dots or the difference in red vs. green dots in the checkerboard. The parameters are governed by the following prior distributions, with fixed hyperparameters:

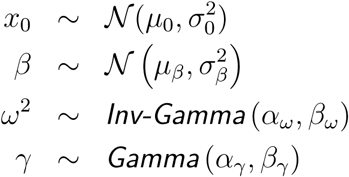

#### Step model

The step model is constructed around three possible firing states: an initial state 0 with constant firing rate *α*_0_ and two other states 1 and 2, with rates *α*_1_ and *α*_2_ respectively, that are associated with the possible decisions in a 2AFC task. Transitions between these states occur instantaneously, and at most one state transition is allowed on each trial (i.e., the firing rate will either remain constant at the initial rate, or change from the initial state to a single other value at some point). The transition on the *j*^th^ trial happens at a time *z_j_* drawn from a negative binomial distribution, and to the state given by *d_j_* ∈ {1, 2}, with probabilities (*ϕ_c_*_(_*_j_*_)_, 1 − *ϕ_c_*_(_*_j_*_)_) that depend on the strength of sensory evidence. If *z_j_* is greater than the trial length, then no step occurs during the trial.

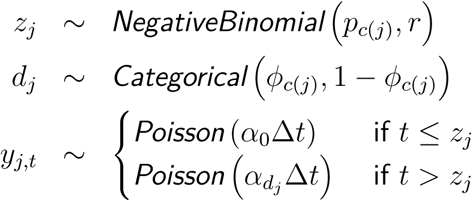

#### Implementation

We used the MATLAB (Mathworks) and CUDA (NVIDIA) code published by Latimer et al. (2015b), with a minor modification to the number of GPU compute threads to accelerate analysis of our PMd, LIP, and hypothetical neuron datasets. We confirmed that our modification did not introduce any inadvertent errors by verifying that our results were consistent with those produced by the original code for two sample neurons. Analyses ran on custom-built computers containing multiple GPUs (9 GPUs in total, 3 in each computer using a mixture of GeForce GTX 980, 980 Ti, Titan Black and Titan Z cards).

### Simulations

Simulations of hypothetical neurons allowed us to study the robustness of model selection, by providing data for which the true generative model was known.

We used the implementation of the DDM “ramp” model (Equations 5–9) provided by Latimer et al. (2015b) as the basis of all the simulations, introducing variations to the core model as described in the following. For simplicity, we generally simulated two sets of trials for each hypothetical neuron: one with a relatively strong positive drift rate, and the other most often with zero drift. While this represented fewer conditions than in the real neural data, it was sufficient to explore the potential brittleness of model selection to underdispersion, non-monotonicity, and parameter variation within a dataset. We also performed some simulations using four different drift rates.

Except where indicated, the default variance of the DDM (*ω*^2^) was 10^−2^. This value was typical of the DDM model parameters fit to our LIP data, as well as those reported by Latimer et al. (2015b). Based on the analysis of the LIP data, we assumed a decision-formation period of 500 ms. When binned using 10 ms bins we obtained a total of 50 time points.

#### Underdispersed responses

We simulated underdispersed DDM responses by first generating time-varying firing rates according to the unchanged DDM (Equations 5–7). Then, rather than generating spike counts from the Poisson distribution assumed during model selection, we instead applied time rescaling to a gamma-interval renewal process to generate spike counts with smaller variance. Specifically, for each trial, we first generated a list of event times *g*_0_, *g*_1_, *g*_2_,… with *g*_0_ = 0 and each interval *g_i_* − *g_i−_*_1_ for *i* ≥ 1 drawn independently from a gamma distribution with mean 1 and shape parameter varying from 1 (corresponding to a Poisson process) to 6 in different simulations. We then set *y_j,t_* to the number of these events that fell between 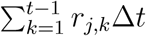 (exclusive) and 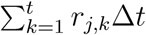 (inclusive). The empty sum in the lower limit when *t* =1 was defined to be 0. When the shape parameter is greater than 1, counts generated in this way will have lower variance than a Poisson process. Indeed, this process generated Fano factors for 10ms-binned counts that ranged from about 0.65 (for shape parameter 6) to 1 (for shape parameter 1, corresponding to Poisson; Supp. Fig. 4A). The trial-to-trial Fano factors for DDM simulations with gamma-interval renewal spiking were somewhat larger as they included variance arising from the latent process.

For these simulations we took *γ*=45 spikes/s, and *x*_0_=0.2, 0.3, 0.35, 0.4, or 0.55. Drift rates for condition 1, ranged from 0.004 to 0.015. For condition 2 drift rate was usually zero and for a small subset of hypothetical neurons was −0.002 to induce a slow decrease in firing rate.

#### Mixed responses

We used two sets of simulations to mimic the complex firing rate profiles found for the PMd neurons.

In the first, we assumed no diffusion noise. Thus the latents of the hypothetical neurons were of the form of a simple deterministic ramp for one condition (condition 1) and a flat level for the other condition (condition 2). Once we obtained latents for each condition, we multiplied them by a non-monotonic function (Fig. 4A), *f*(*t*), that was defined as follows.

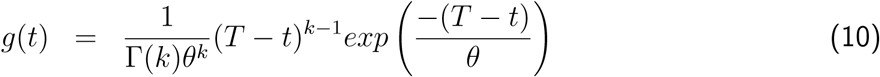

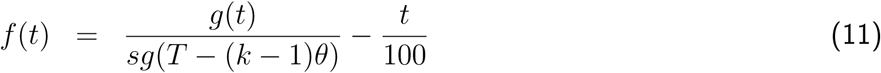

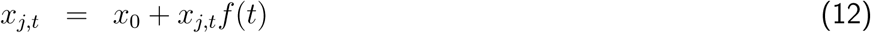

We assumed T=55, k=5, s=1.1, and *θ*=4 and that the time variable *t* was 50 time points long starting from t=1.

In the second set of simulated non-monotonic neurons, we assumed both drift and diffusion (again with *ω*^2^ = 10^−2^). Once we had simulated initial latents from the DDM, we then again used the function defined in Eqn. 11 above to create a non-monotonic firing rate profile. These latents were then converted into spike counts after being mapped into firing rates using equation 9.

We also assumed for some simulations of hypothetical neurons, four stimulus levels, instead of just two stimulus levels.

#### Non-stationary parameters

To explore brittleness in the face of non-stationarity we generated data from a DDM in which diffusion noise varied with condition. The common parameters *γ*=45spikes/s and *x*_0_ =0.4 were shared by all simulations. We varied *ω*^2^ and *β_c_* for the two choices. We again assumed non-zero drift rates for trials from condition 1 (range from 0.0035 – 0.0190) and a 0 drift rate for condition 2. Condition 1 trials were simulated with a diffusion variance *ω*^2^ = 10^−2^. The diffusion variance on condition 2 trials was set either one or two orders of magnitude smaller: *ω*^2^ = 10^−3^ or 10^−4^. Given these parameters, spike counts were generated using the DDM model (Equations 5–9).

#### Hybrid model

Finally, we considered hypothetical neurons that ramp under one condition (corresponding to PREF choices) but step in another. For simulation of PREF-choice firing rates, we assumed the following parameters for the DDM: *γ* = 50, *x*_0_ =0.25, *ω*^2^ = 10^−2^, and *β* ranging from 0.0045 to 0.0160. For the NONPREF-choice firing rates, we assumed a step model with negative binomial parameters *p* =0.92, *r* =2; *ϕ* =0.9; *α*_0_ = *α*_2_ = 12.5 spikes/s and *α*_1_ = 15 spikes/s or 10 spikes/s.

### Deviance Information Criterion for model selection

Latimer et al. (2015b) performed model selection using the deviance information criterion (DIC), which provides a Bayesian estimate of the divergence error of a model (Gelman et al., 2014; Spiegelhalter et al., 2002). Despite its Bayesian formulation, DIC is closer in spirit to Akaike’s (1974) Information Criterion (AIC) than to the Bayesian Information Criterion (BIC; Schwarz, 1978) in that it seeks to find the closer of the models to the data rather than choosing the one most likely to be correct (an effort of debatable utility when the answer is almost certainly “neither”). Unlike AIC, DIC incorporates prior information and provides an estimate that should be useful outside an asymptotic limit. It is well-suited for use with Markov chain Monte-Carlo fitting methods, which draw samples from the posterior over the model parameters even when the exact posterior density cannot be computed. Nonetheless, it is open to criticism (Spiegelhalter et al., 2014). We employed it here to maintain compatibility with the earlier study, and our central point concerned the interpretation of model selection methods in general rather than any particular criterion.

The DIC for a model 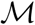 with parameters 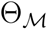 is defined as:

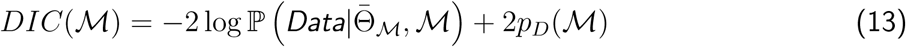

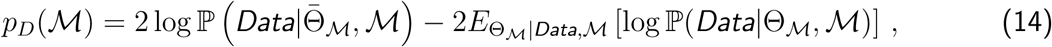

where *Data* represents the available observations and 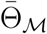 denotes the posterior mean of the model parameters given these data.

The first term in Equation 13 is the deviance: twice the negative log-likelihood, usually measured relative to a baseline model although we have omitted that term here as it does not affect the outcome of model selection. A lower value indicates a better fit. The quantity *p_D_* is an estimate of the discrepancy between the deviance of the mean parameters and the expected posterior of the “true deviance”. It acts as a form of data-dependent model complexity penalty.

Both the mean parameters 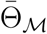 and the expectation in the second term of *p_D_* (Equation 14) can be estimated by taking the corresponding empirical averages over Monte-Carlo samples from the posterior distributions 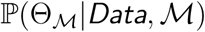. Evaluaton of the likelihoods also require a numerical estimate of the implicit integral over the latent parameters. For the DDM, we used the code from (Latimer et al., 2015b) that computed Monte-Carlo estimates with 300 sample trajectories for each trial and each parameter value. For the step model, we truncated the possible step times to a maximum of 1500ms and evaluated the likelihood using a grid-based numerical integral.

#### DIC score

For model selection, DIC is computed separately for the DDM and step models. Larger DIC values mean that the model is a poorer fit to the data. We compared the models using a relative DIC “score”, which was the difference between the DIC values for the two models:

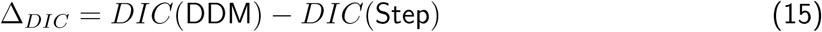

A DIC score > 0 suggests that the step model describes single-trial firing rates better than the DDM, and a DIC score < 0 implies the opposite.

#### DIC contributions from PREF and NONPREF

Under some circumstances, it is useful to understand the contribution of different types of trials to the DIC value for a given model. In this study the data were binned spike counts recorded in independent trials. Thus the log probabilities in Equations 13–14 can be written as sums of single-trial probabilities, giving:

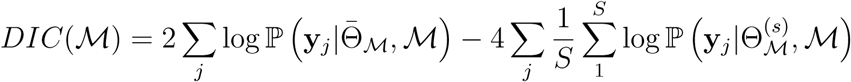

where **y***_j_* represents the binned spike counts on trial j. Thus, we can define a “single-trial DIC contribution” for the j^th^ trial by

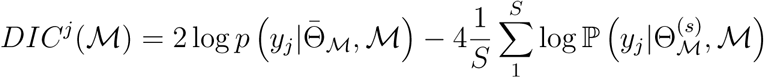

Note however that the mean parameter 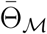 and samples 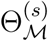 are still computed using all trials so this is not a DIC value in itself – rather it measures a contribution to the overall value obtained from this trial. Such single trial contributions can be summed over subsets of trials—for example those corresponding to PREF or NONPREF choices—to examine the net contributions from these different trial types; and the difference of these contributions assessed for the two different models provides a useful measure of relative impact of different trial types on ranking. For example

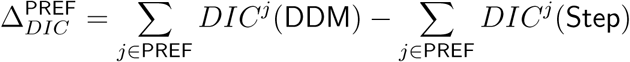

Systematic differences in these contributions across different trial types may suggest a fundamental incompatibility in both models.

## Supplementary Figures

**Supplementary Figure 1:**
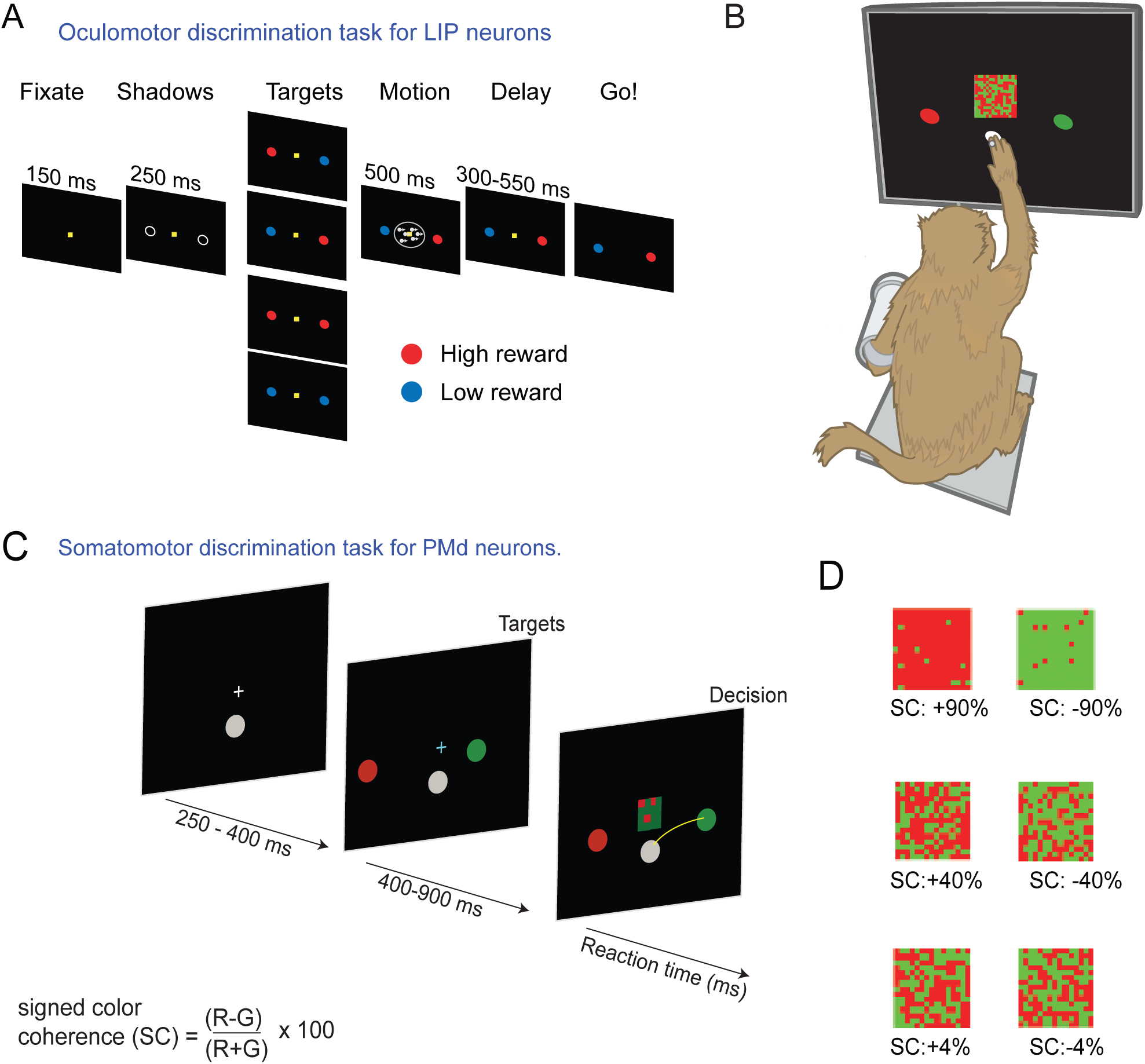
Tasks performed by monkeys for recording of LIP neurons and PMd neurons. **A:** Sequence of events comprising a typical trial in the oculomotor fixed-duration discrimination task used in Rorie et al. (2010). Figure adapted from Figure 1 of Rorie et al. (2010). From left to right, trials begin with the onset of a fixation point. Shortly after the monkey fixates its gaze on the fixation point, two saccade targets appear and then change color indicating the magnitude of the reward available for correctly choosing that target. A blue color target indicates a low magnitude (L) reward, while a red color target indicates a high magnitude (H) reward. The four reward conditions are depicted vertically—HL, LH, HH, LL, from top to bottom. The visual motion stimulus is centered on the fixation point. Following offset of the motion stimulus, the animals wait for a short randomized delay period (300 – 550 ms), and then the fixation point extinguishes and the animal is free to make its choice. A successful trial is rewarded with a drop of juice. **B:** An illustration of the setup for the behavioral task used for recording decision-related activity in the reaction time visual discrimination task used for PMd. We gently restrained the arm the monkey was not using with a plastic tube and cloth sling. We tracked a reflective IR bead taped on the middle digit of the hand to mimic a touch screen and to provide an estimate of instantaneous arm position. Eye position was tracked using an infra-red reflective mirror placed in front of the monkey’s nose. **C:** Time line of the somatomotor reaction time discrimination task used for recording of PMd data in Chandrasekaran et al. (2017). From left to right, trials begin with the onset of a central hold target and a fixation cross. Shortly after the monkey places his hand on the central hold and fixates on the cross, two reach targets appear on either side of the central target. The targets are red and green in color. On some trials the left target is red and the right target is green and vice versa. After a brief holding period, a central static checkerboard composed of red and green squares appears. The task is a reaction time task and thus the monkey is free to initiate a reach whenever he feels he has sufficient evidence to report his decision. **D:** Examples of different stimulus ambiguities used in the experiment parameterized by the color coherence of the Checkerboard cue defined as SC=100×(R-G)/(R+G). Positive values of signed color coherence denote more red than green squares and vice-versa.

**Supplementary Figure 2:**
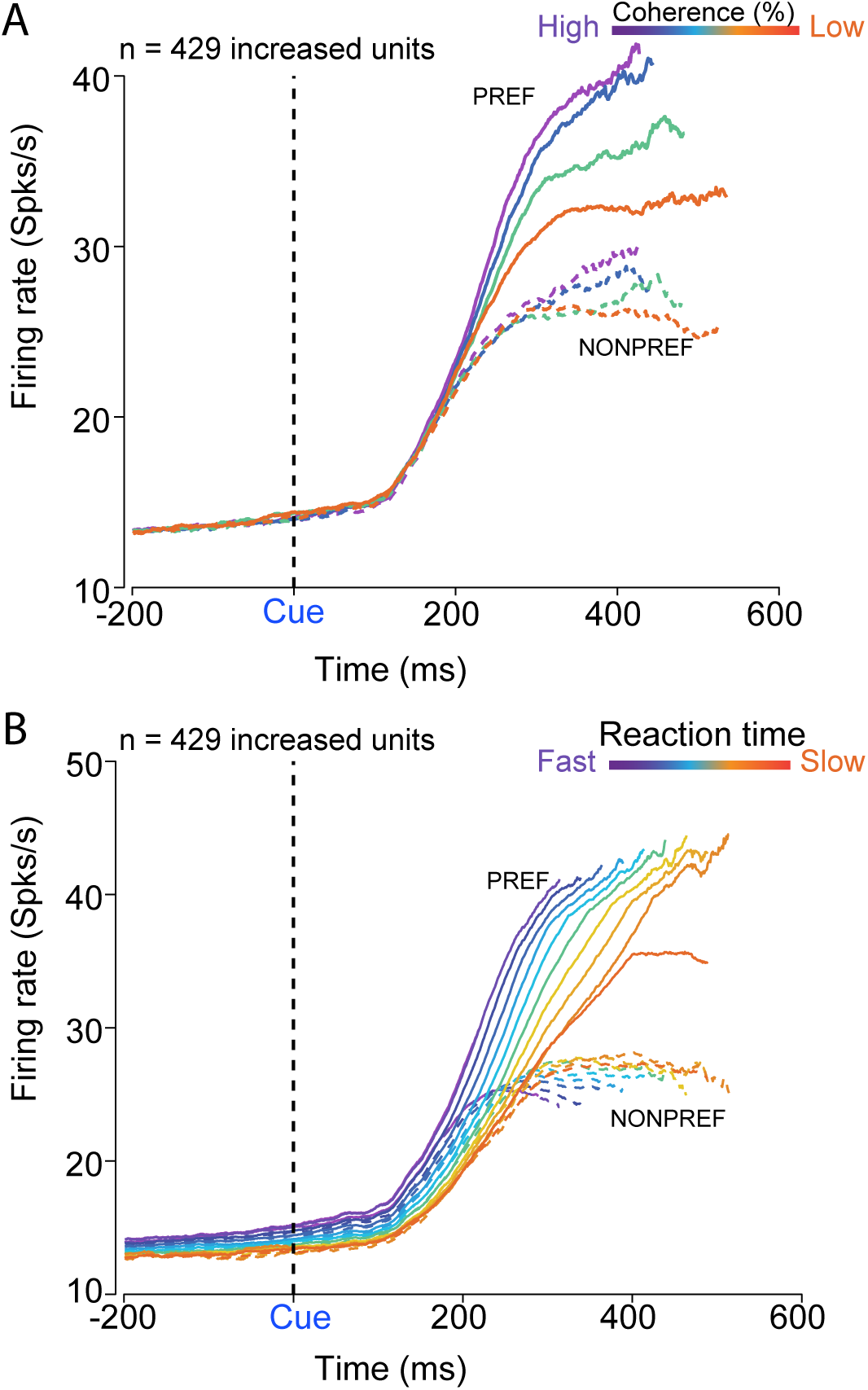
Average firing rate of increased PMd units aligned to checker-board onset, organized by reaches into the PREF or NONPREF direction, and sorted by either coherence or reaction time. **A**: Average firing rate across all 429 increased units considered in this study sorted by the coherence of the checkerboard and the choice made by the animal. Colors label coherence conditions from easy (purple) to difficult (orange). Solid lines show movements to the PREF-direction; dashed lines show the NONPREF-direction. firing rate traces are obtained using 50ms causal boxcar filters, and, in the CUE-aligned upper panels, truncated at the center point of the reaction-time range for each coherence. **B**: Average firing rate across all 429 increased units considered in this study sorted by the reaction time of the animals and the choice made by the animals. Solid lines depict reaches for the PREF direction; dashed lines depict the NONPREF direction. Colors denote different reaction time bins.

**Supplementary Figure 3:**
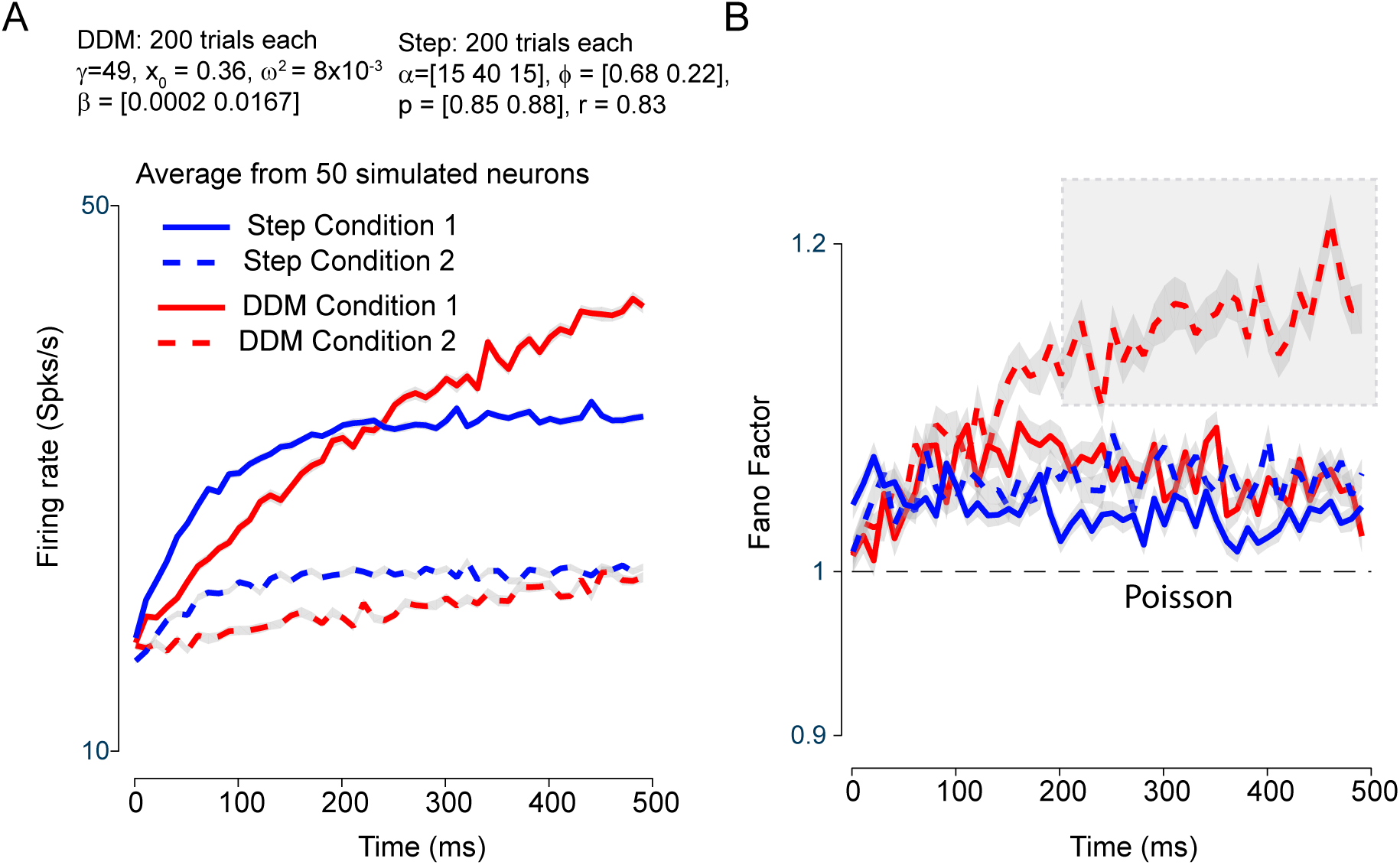
Both step model and DDM assume that Fano factor is super-Poisson (or overdispersed) **A:** Average firing rate for a condition with strong drift rates or large firing rate step (solid lines) or modest drift rates or steps (dashed lines) directions of 50 hypothetical neurons simulated either from the DDM (red line) or the step model (blue line) variant described in Latimer et al. (2015b). The magnitudes of the various parameters for the DDM and the step model were chosen so that rates of the hypothetical step neurons and hypothetical DDM neurons roughly matched. **B:** Average Fano Factor for the same neurons shown in A. Figure conventions as in A. Fano Factor for both the step model and the DDM are super-Poisson. We also note that Fano factor for modest firing rates, which usually emerges from small drift rates, steadily increases with time for neurons simulated from the DDM as defined by Latimer et al. (2015b). The shaded grey rectangle is to draw the reader to the rapid increase in Fano factor for modest drift rates when simulated from the DDM.

**Supplementary Figure 4:**
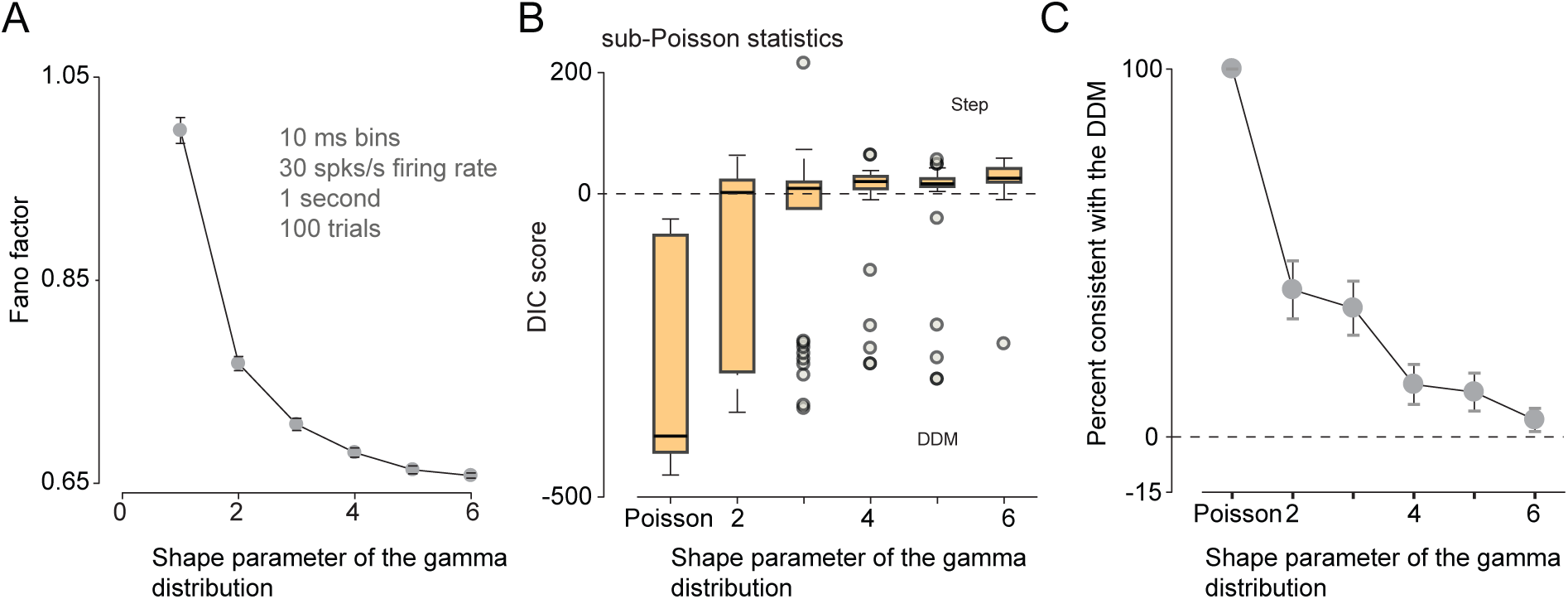
Sub-Poisson firing rates impact results of model selection even with multiple coherences. **A**: Shows the simulated Fano factor for the gamma-interval renewal process. Data are averages over 100 trials simulated at 30 Hz for 1 second each, binned in 10 ms bins. As the shape parameter for the gamma distribution increases, the Fano factor decreases from 1. **B**: Box plot of DIC score distribution for DDM simulations with 4 different drift rates and with gamma-interval renewal process spiking, as a function of the shape parameter of the gamma-interval distribution. Higher values of shape parameter lead to increasing spiking regularity and thus sub-Poisson Fano factors. DIC scores for simulated neurons with DDM dynamics were increasingly identified as consistent with the step model when the firing rates became more sub-Poisson. Large outlier DIC scores (grey dots) sometimes support the DDM for these neurons, but the DIC scores for a majority of these underdispersed neurons are consistent with the step model. Each box plot is estimated from the DIC scores of 45 neurons. Only neurons with robust firing rate modulation were considered for this plot and panel B. **C**: Fraction of neurons consistent with the DDM decreases as the shape parameter of the gamma-interval renewal process increases. A neuron was considered consistent with the DDM if DIC was less than 0 for the purposes of this study. Note that neurons quickly become inconsistent with the DDM even with minor levels of underdispersion.

**Supplementary Figure 5:**
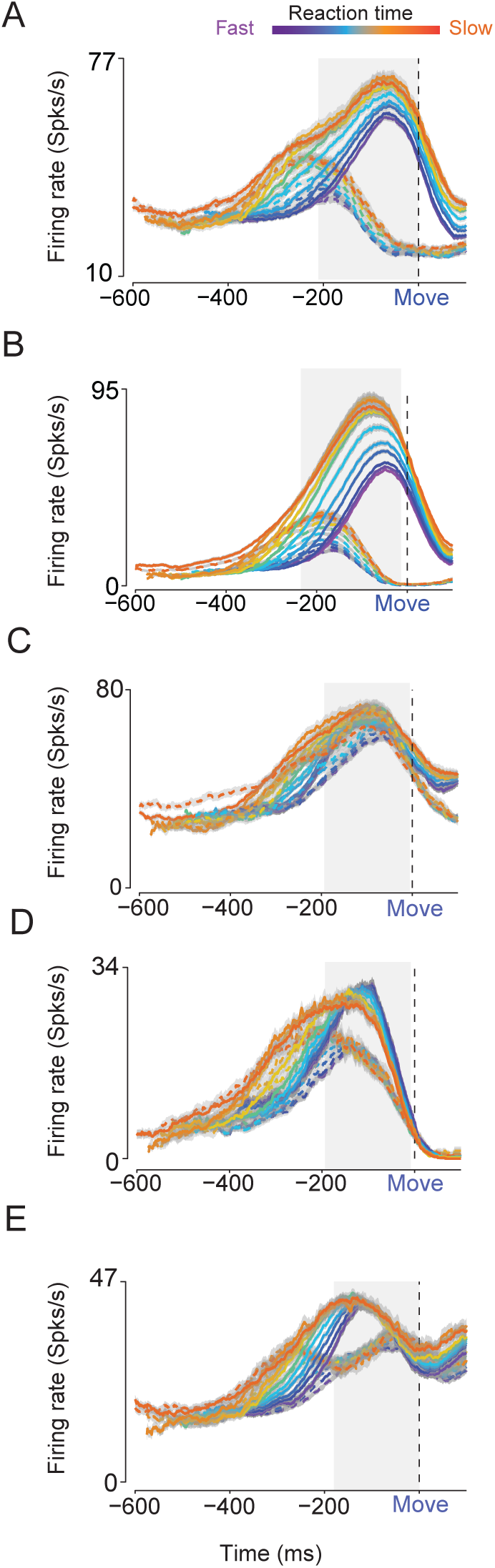
Additional examples of PMd units that show non monotonicity in their firing rates as a function of time. **A-E**: PREF and NONPREF firing rates for five other PMd units aligned to movement onset. Solid lines show PREF firing rates, dashed lines show NONPREF choice firing rates. Colors from purple to orange show different reaction time bins. Dashed black line denotes movement onset. Gray shading is provided to orient the eye to the non-monotonicity in the firing rates.

**Supplementary Figure 6:**
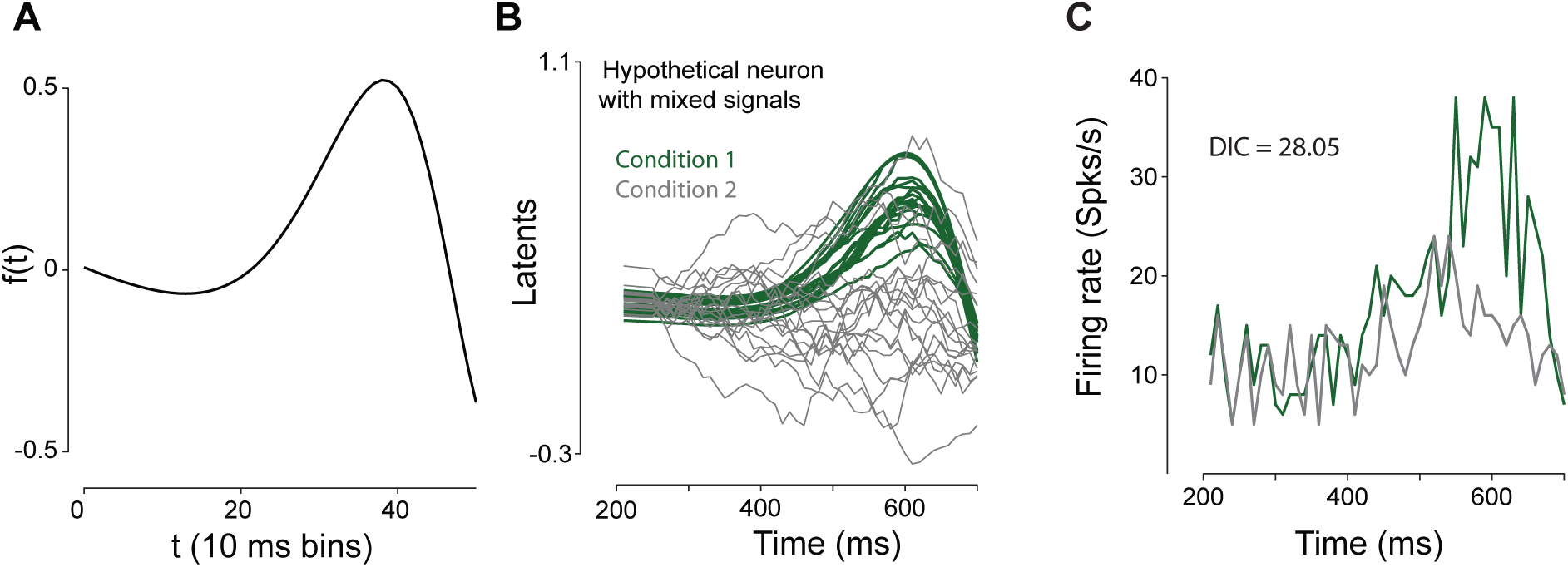
Another example of a hypothetical PMd neuron with non-monotonic firing rate profile being identified as consistent with the step model. **A**: A plot of f(t), the non-monotonic function (see Eqn. 11) used for simulation of hypothetical neurons that mimic the responses of PMd neurons. f(t) is typically multiplied by *x_j_*(*t*) to give rise to nonmonotonic latent profiles that are in turn converted into firing rates. **B**: Latent dynamics of another example of a hypothetical neuron meant to mimic the firing rate patterns observed in PMd. The existence of many more traces in A compared to Fig. 4C are due to the latents initially being generated from a DDM and then multiplied by a non-monotonic profile. Green traces reflect one condition and gray traces reflect another condition. **C**: Trial-averaged firing rates of the hypothetical neuron shown in A. For each trial spike trains were generated using a Poisson model output process with latent dynamics given in A. Firing rates were obtained by averaging over trials for each choice. Note that no steps were involved in the simulation but the model selection method returns DIC scores that identify the neuron as better described by the step model.

## Author Contributions

CC collected the PMd data with input from KVS, WTN and DP, wrote the code, constructed simulations, analyzed all data, and made figures. JSM and MS constructed simulations and also developed analytical models to guide insight. CC, KVS, and MS wrote initial drafts of the paper. LIP data were collected in the lab of WTN by Dr. Alan Rorie. All authors participated in the development of the figures, interpreted results of data analyses, suggested simulations and analyses, and revised the final drafts of the paper.

## Acknowledgments

The work was supported by the following grants:

- CC was supported by an NIH/NINDS K99/R00 grant NS092972 and the Howard Hughes Medical Institute.
- JSM was supported by the Gatsby Charitable Foundation
- DP was supported by the Champalimaud Foundation, Portugal and Howard Hughes Medical Institute
- BN was supported by the Howard Hughes Medical Institute.
- KVS was supported by a Defense Advanced Research Projects Agency NeuroFAST award from BTO #W911NF-14-2-0013 and the Howard Hughes Medical Institute.
- MS was supported by the Gatsby Charitable Foundation and the Simons Foundation (SCGB 323228, 543039).

We thank Tatiana Engel for helpful suggestions and also advice on implementing time-rescaled renewal processes.

